# Environmental effects on genetic variance are likely to constrain adaptation in novel environments

**DOI:** 10.1101/2021.02.08.430333

**Authors:** Greg M. Walter, Keyne Monro, Delia Terranova, Enrico la Spina, Maria Majorana, Giuseppe Pepe, James Clark, Salvatore Cozzolino, Antonia Cristaudo, Simon J. Hiscock, Jon R. Bridle

## Abstract

Adaptive plasticity allows populations to cope with environmental variation but is expected to fail as conditions become unfamiliar. In novel conditions, populations may instead rely on rapid adaptation to increase fitness and avoid extinction. Adaptation should be fastest when both plasticity and selection occur in directions of the multivariate phenotype that contain abundant genetic variation. However, tests of this prediction from field experiments are rare. Here, we quantify how additive genetic variance in a multivariate phenotype changes across an elevational gradient, and test whether plasticity and selection align with genetic variation. We do so using two closely related, but ecologically distinct, sister species of Sicilian daisy (*Senecio,* Asteraceae) adapted to high and low elevations on Mount Etna. Using a paternal half-sibling breeding design, we generated and then reciprocally planted c.19,000 seeds of both species, across an elevational gradient spanning each species’ native elevation, and then quantified mortality and five leaf traits of emergent seedlings. We found that genetic variance in leaf traits changed more across elevations than between species. The high-elevation species at novel lower elevations showed changes in the distribution of genetic variance among the leaf traits, which reduced the amount of genetic variance in the directions of selection and the native phenotype. By contrast, the low-elevation species mainly showed changes in the amount of genetic variance at the novel high elevation, and genetic variance was concentrated in the direction of the native phenotype. For both species, leaf trait plasticity across elevations was in a direction of the multivariate phenotype that contained a moderate amount of genetic variance. Together, these data suggest that where plasticity is adaptive, selection on genetic variance for an initially plastic response could promote adaptation. However, large environmental effects on genetic variance are likely to reduce adaptive potential in novel environments.

## Introduction

Populations facing rapid environmental change must cope with increasingly novel environments if they are to persist. Plastic responses to different environments can aid persistence by allowing populations to rapidly change their mean phenotypes to track changes in phenotypic optima, maintaining fitness in each environment (Via et al. 1995; Charmantier et al. 2008). However, plasticity should only evolve to track environments that populations regularly experience (Hermisson and Wagner 2004; Chevin et al. 2010; Ashander et al. 2016), and is unlikely to maintain fitness in novel environments because plastic responses have not evolved to suit those environments (Palacio-López et al. 2015; Acasuso-Rivero et al. 2019). When plasticity fails to maintain fitness, genotypes can vary in the extent to which they suffer reduced fitness, increasing the potential for populations to recover fitness when selection on ecologically important traits leads to adaptation (Shaw and Shaw 2014; Walter et al. 2023). Our understanding of the potential for adaptation in novel environments is limited because field experiments that measure plasticity, selection and genetic variation along ecological gradients are rare.

For adaptation to occur, selection must operate on genetic variation in multiple traits that combine to form a multivariate phenotype (Blows 2007). However, pleiotropy and linkage often create genetic correlations among traits, meaning that any changes to one trait will enact changes in other traits that are genetically correlated. The additive genetic variance-covariance matrix (**G**) summarises how genetic variation is distributed among a given set of traits (**Fig. 1a**; Lande 1979; Walsh and Blows 2009). The diagonal of **G** captures genetic variation in each trait, while the off-diagonal of **G** measures the covariation between each pair of traits due to shared genetic variation (Steppan et al. 2002). Stronger covariances between traits concentrate the total variance in **G** into certain trait combinations (**Fig. 1a**). These are often found by decomposing **G** into independent axes, akin to principal components, representing directions in which phenotypes vary genetically (Lande 1979; Walsh and Blows 2009). The primary axis of **G**, known as ***g***_max_, is the direction of the phenotype in which genetic variation is most abundant and the direction in which phenotypic evolution is expected to occur (Schluter 1996; McGlothlin et al. 2018; Costa e Silva et al. 2020; Zu et al. 2020; Walter 2023).

**Fig. 1.**
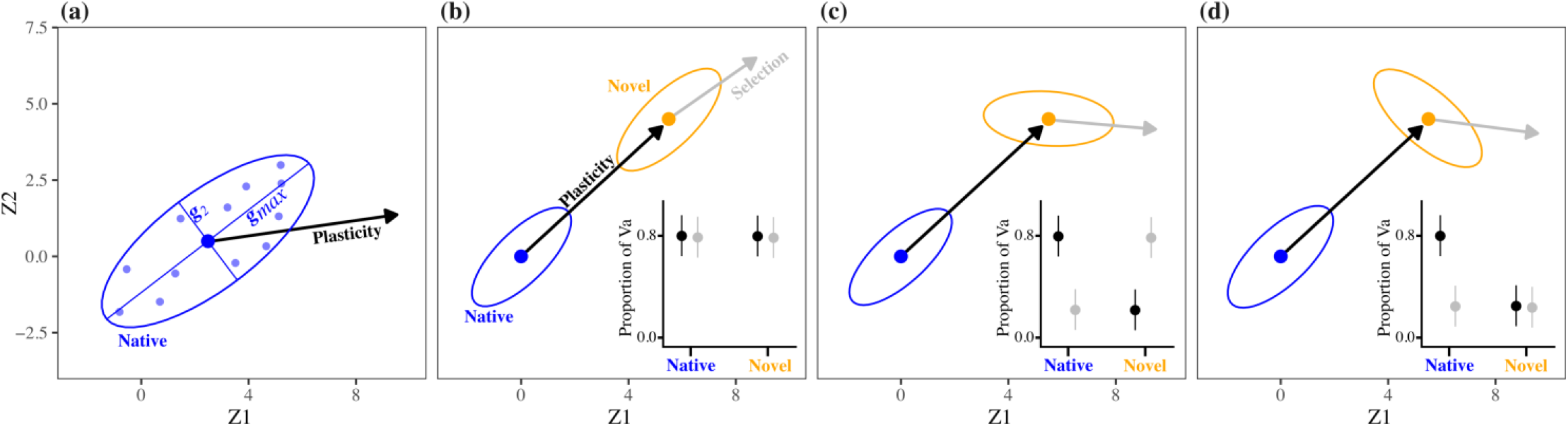
**(a)** A genetic correlation between two traits (Z1 and Z2) concentrates genetic variation in the traits (summarised by the additive genetic variance-covariance matrix, **G**) along axes defined by trait combinations. The ellipse shows genetic variation in the traits as a cloud of genotypes (small circles) in multivariate space with ***g***_max_ representing the direction in which both traits vary most genetically. In this case, genotypes with larger values of Z1 also have larger values of Z2. Plasticity is the change in trait mean (large circle) in response to a change in environment. **(b-d)** Comparing plasticity and selection with genetic variation for Z1 and Z2 in a population at its native elevation (blue) and a novel elevation (orange). Again, ellipses show genetic variation in traits and solid circles within ellipses are trait means. Black arrows are changes in trait means across environments due to plasticity, and grey arrows are the directions of selection at the away site. Insets show the proportions of genetic variation in traits in the direction of plasticity and selection in each environment. Adaptation will be fastest if both plasticity and selection are in a direction with a large amount of genetic variance in the traits (i.e., ***g***_max_), as in **(b)**. However, genotype-by-environment interactions (G×E) mean that genotypes vary in plasticity, which can change genetic variation in traits across environments, as in **(c-d)**. In the novel environment, if plasticity creates phenotypes that differ to those favoured by selection, then plasticity will be nonadaptive. In this case, for rapid adaptation to be possible, G×E interactions would need to increase genetic variation in the direction of selection, as in **(c)**. However, adaptation in the novel environment will be constrained if G×E reduces the amount of genetic variation in the direction of selection in that environment, as in **(d)**.

In novel environments, plasticity will be at least partially adaptive if it moves a population’s mean phenotype closer to its new optimum (Ghalambor et al. 2007; Lande 2009). Evidence from diverse taxa also suggests that plasticity, adaptive or otherwise, can be biased towards the direction of ***g***_max_ (Lind et al. 2015; Noble et al. 2019). Together, these observations predict that adaptation in novel environments will be fastest if plasticity, selection, and genetic variation in the same phenotype are all aligned (**Fig. 1b**; Lande 2009; Levis and Pfennig 2016). However, if plasticity is non-adaptive, which is common (Acasuso-Rivero et al. 2019), then its alignment with ***g***_max_ could deflect evolution away from the direction of selection and thereby constrain adaptation to the new environment. Testing these predictions requires field experiments that use natural populations to test whether plastic responses to novel environments are (1) adaptive and (2) produce phenotypes with abundant genetic variation.

The availability of genetic variation can depend on whether genotypes vary in plasticity, known as genotype-by-environment interactions (G×E) (Josephs 2018). The presence of G×E in a set of traits means that additive genetic variation for the traits and/or covariation between them (summarised in **G**) changes with the environment (Sgrò and Hoffmann 2004; Wood and Brodie III 2015). This can aid adaptation to novel environments when G×E causes genetic variation to increase in the phenotypes favoured by selection (**Fig. 1c**). Conversely, G×E that reduces genetic variation in phenotypes under selection should constrain adaptation to novel environments (**Fig. 1d**; Chevin 2013). Theory supports the first of these scenarios, predicting that changes in **G** in new environments aid adaptation by aligning the directions of ***g***_max_ and selection (Draghi and Whitlock 2012; Chevin 2013; Lind et al. 2015). To our knowledge, however, no field experiments have quantified plasticity, selection, and genetic variation along environmental gradients as current ecological margins are exceeded. Consequently, we do not yet know whether changes in genetic variance across environments increase the adaptive potential of populations when faced with novel conditions.

We planted seeds of two closely related, but ecologically contrasting, sister species of Sicilian daisy (*Senecio*, Asteraceae) across an elevational gradient on Mt. Etna (**Fig. 2a-b**). Both species are obligate outcrossers that share generalist insect pollinators (Walter et al. 2020a). *Senecio aethnensis* is a perennial with entire glaucous leaves that is endemic to lava flows >2,000 metres above sea level (m.a.s.l.) on Mt. Etna, where individuals grow back each spring after snow cover in winter. By contrast, *S. chrysanthemifolius* is a short-lived perennial with dissected leaves that occupies disturbed habitats (e.g., roadsides and vineyards) at 500-1,000m.a.s.l. on Mt. Etna and across Sicily more broadly. Despite its narrower geographical distribution, *S. aethnensis* has greater genetic diversity than *S. chrysanthemifolius,* suggesting that *S. aethnensis* derives from a larger ancestral population (Chapman et al. 2013). However, *S. chrysanthemifolius* shows greater adaptive phenotypic plasticity across elevation than *S. aethnensis* (Walter et al. 2022a).

**Fig. 2.**
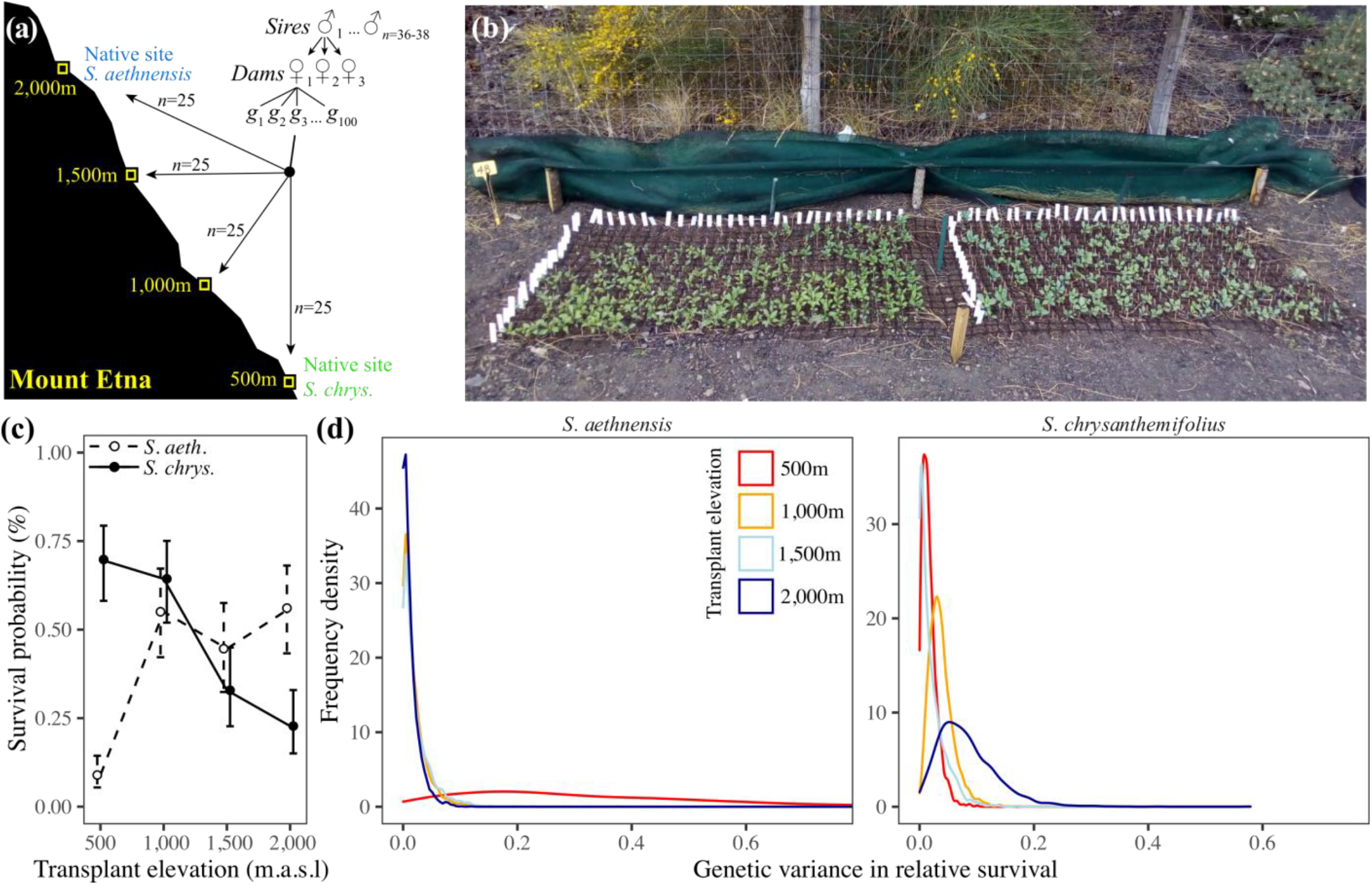
The experimental design. **(a)** For two Etnean *Senecio* species, we mated each of 36-38 sires to each of 3 dams, and collected 100 seeds from each mating (family). We planted 25 seeds from each family at each of four elevations representing the native elevation of each species and two intermediate elevations. **(b)** Photo of an experimental block at 2,000m, 8 weeks after seeds were sowed (*S. chrysanthemifolius* on left). Panels **c-d** show analyses of survival data from this experiment, taken from Walter et al. (2022b). **(c)** Mean survival probability after summer (±1 SE) for species across the elevation gradient. **(d)** Posterior distributions for estimates of genetic variance in survival, which was greater for both species at the novel elevation.

To estimate additive genetic variance in phenotypes and fitness, we used a paternal half-sibling breeding design to produce seeds for c.100 families per species. We then reciprocally planted seeds from each family across an elevational gradient spanning the native ranges of both species and two intermediate elevations. We tracked the survival of all seedlings that emerged, and measured five ecologically-important leaf traits that are known to be plastic (Walter et al. 2022a), correlated with fitness (Walter et al. 2023), and associated with adaptive divergence in other *Senecio* species (Richards et al. 2019; Walter et al. 2020b). Previous analyses of survival data from this experiment show that these two species are adapted to their native environments (**Fig. 2c**) and show increased additive genetic variance in survival that improves their adaptive potential when planted in novel environments (**Fig. 2d**) (Walter et al. 2022b).

Here, we extend these results by estimating the additive genetic variance-covariance matrix (**G**) for the five leaf traits measured in each species at each elevation, allowing us to compare plasticity, selection and genetic variation in leaf traits along a natural ecological gradient. We first compare **G** across elevations to test whether G×E causes genetic variation in leaf traits to change in response to this natural environmental gradient, predicting that **G** would differ the most between native and more novel elevations. We then quantify how much genetic variance lies in the direction of plasticity and selection at novel elevations, predicting that G×E would aid adaptation in novel environments if there is abundant genetic variance in the direction of leaf phenotypes favoured by selection.

## Methods and materials

### Breeding design and field experiment

We briefly describe the field experiment and refer readers to Walter et al. (2022b) for more detail. We collected cuttings from c.80 individuals per species growing naturally at 2,000-2,600m.a.s.l for *S. aethnensis* and 526-790m.a.s.l for *S. chrysanthemifolius.* (**Table S1** and **Fig. S1)**. We propagated one cutting per field individual in the glasshouse, and randomly assigned the individual as a sire (pollen donor) or dam (pollen receiver). We randomly mated three sires to three dams in full-factorial 3×3 blocks, completing twelve blocks for *S. aethnensis* (*n*=36 sires, *n=*35 dams, *n*=94 full-sibling families) and 13 blocks for *S. chrysanthemifolius* (*n=*38 sires, *n*=38 dams, *n*=108 full-sibling families, with two sires and dams in the last block). We then planted 100 seeds from each family on Mt. Etna at four elevations, including the native elevations of both species (500m and 2,000m) and two intermediate elevations (1,000m and 1,500m). Vagrant individuals of both species are occasionally found at the intermediate elevations, suggesting the potential to expand their ranges beyond their current distributions.

At each site, we planted 25 seeds randomised into five experimental blocks (*S. aethnensis n*=432 seeds/block, *n=*2,160 seeds/site; *S. chrysanthemifolius n=*540 seeds/block, *n=*2,700 seeds/site; total N=19,232 seeds). To prepare each experimental block, we cleared the ground of plant matter and placed a plastic grid (4cm-square cells) on the ground. We attached each seed to the middle of a toothpick using non-drip super glue and then pushed each toothpick into the soil in each grid cell so that the seed sat 1-2mm below the soil surface. To replicate natural germination conditions, we suspended 90% shade-cloth 20cm above each block and kept the seeds moist until germination ceased (2-3 weeks). We then replaced the 90% shade-cloth with 40% shade-cloth to replicate shade that naturally-growing plants are often found under, which maintained a strong environmental gradient across elevations without exposing plants to extreme conditions. We recorded seedling emergence, survival and establishment (whether seedlings produced ten leaves). The experiment ended in January 2020 when mortality stabilised (**Fig. S2**) and plants started growing into each other, increasing competition. This precluded further data collection, including reproductive traits.

### Quantifying leaf traits

When more than 80% of plants had produced ten leaves at each transplant elevation, we collected the 5^th^ and 6^th^ leaves (from the base of plant) to quantify leaf morphology and pigment content (N=6,454 plants). For leaf morphology, we scanned the leaves (Canoscan 9000F) and used *Lamina* (Bylesjo et al. 2008) to quantify leaf complexity 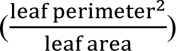, width of leaf indents (mm) and number of leaf indents standardised by perimeter (indents/mm). We then weighed all leaves for each plant and calculated specific leaf area (SLA) as 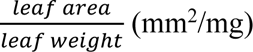. We used a Dualex instrument (Force-A, France) to measure the flavonol content of each leaf in spectral reflectance.

Prior to analyses, we mean standardised each trait to aid the comparison of traits measured on different scales (Hansen and Houle 2008). This also estimates the mean-standardised genetic variance (evolvability) of each trait, as appropriate for comparison with changes in trait means (plasticity) across elevations. We used R (v.3.6.1; R Core Team 2021) for all analyses.

### Quantifying plasticity as elevational changes in multivariate phenotype

To quantify species differences in leaf plasticity across elevations, we used a multivariate analysis of variance (MANOVA) with the five leaf traits as the multivariate response variable, and with elevation, species, and their interaction as fixed effects. Block within elevation, and family within species, were error terms for species and elevation respectively. To visualise differences in plasticity, we estimated the D-matrix of differences in mean multivariate phenotype and calculated scores for the first two axes of **D**. Methods for constructing **D** are presented in **Methods S1**.

### Estimating additive genetic variation in leaf traits

To estimate the additive genetic variance-covariance matrix (**G**) for the five leaf traits measured in each species at each elevation, we used the package *MCMCglmm* (Hadfield 2010) to apply the linear mixed model

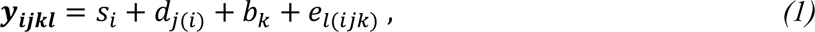

where leaf traits are the multivariate response variable (***y_ijkl_***), *s*_*i*_ is the *i*th sire, *d_j(i)_* the *j*th dam nested within sire, *b_k_* the *k*th block, and *e*_*l*(*ijk*)_ the residual variation. We applied equation 1 separately to the data for each species at each elevation (*n*=8). We used chains with burn-ins of 150,000 iterations and thinning intervals of 1,500 iterations, saving 2,000 thinned iterations (MCMC samples) as the posterior distributions for all estimates. We confirmed model convergence by checking that chains mixed sufficiently well, that autocorrelations among samples were below 0.05, and that our parameter-expanded prior was uninformative (Hadfield 2010). For each model, we constructed **G** with the sire variances and covariances, which represent one-quarter of the additive genetic variation in leaf traits (Lynch and Walsh 1998).

Since *MCMCglmm* constrains variance estimates to be positive, we tested the significance of estimates in **G** by comparing them with suitable null distributions created by randomising offspring among sires and re-applying the model to randomised data. To maintain differences among blocks, we randomised offspring within each block separately. We conducted 1,000 randomisations for each observed **G** and concluded that observed estimates in **G** were significant if posterior means exceeded those from null distributions.

Due to slower growth at higher elevations, many plants died before they could be measured, which could influence our estimates of genetic variance (**Fig. S2**). Nevertheless, we measured multiple offspring from all sires and more than 90% of full-sibling families (4.5-15.6 offspring per family on average), meaning that our estimates of **G** are based on the entire pedigree and large sample sizes (*n*=482-1,683 individuals) (**Table S2**).

### Comparing additive genetic variation across elevations and species

To quantify differences in genetic variation in leaf traits across elevations and species, we used eigenanalysis to decompose each **G** into independent axes (eigenvectors) defined by linear combinations of traits that represent directions in which leaf phenotypes vary genetically. As such, eigenvectors describe the orientation of genetic variation in leaf phenotypes expressed by each species at each elevation, and have eigenvalues describing the amounts of genetic variation in those phenotypes. We used these descriptors to characterise genetic variation in leaf phenotypes and how genetic variation changes across elevations.

For each G-matrix, we used the total amount of genetic variation in leaf traits to describe its size, the distribution of eigenvalues to describe its shape (more elliptical if variation is more condensed towards certain phenotypes), and eigenvectors to describe its orientation. To compare the orientation of genetic variation between elevations, we calculated the angle between ***g***_max_ estimated at the native elevation and ***g***_max_ estimated at each of the other elevations using

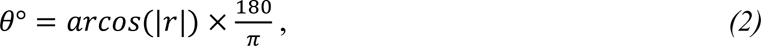

where *r* is the correlation between eigenvectors and *θ*° is the angle between them (Lind et al. 2015; Berdal and Dochtermann 2019).

We then used a covariance tensor approach to formally compare **G** matrices within a single framework. Briefly, the covariance tensor is an eigenanalysis of the S-matrix, which contains the variances and covariances of the individual elements in **G** across our eight matrices. Decomposing **S** therefore provides (after re-arrangement and scaling) a set of independent axes (eigentensors) describing differences in **G** across elevations and species. For more details, see **Fig. S3** and Hine et al. (2009); Aguirre et al. (2014); Walter et al. (2018).

To test the significance of observed eigentensors, we compared them to suitable null distributions created by re-applying equation 1 to data re-constructed from randomised breeding values, simulating changes in **G** due only to random sampling (see supplementary code; Morrissey et al. 2019; Walter 2023). If observed eigentensors described larger differences in **G** compared to null eigentensors, we concluded that genetic variance for leaf traits differed significantly across elevations and/or species. To identify how each original **G** contributed to such differences, we calculated matrix coordinates. Like principal components scores, coordinates describe correlations between eigentensors and the original matrices. Matrices with larger scores contribute more to overall differences described by an eigentensor.

### Comparing plasticity and selection with genetic variation in leaf traits across elevations

#### Does plasticity occur in directions of phenotype with abundant genetic variation?

For each species, we quantified multivariate plasticity in leaf traits between the native elevation and the other elevations. We first calculated the mean of all five leaf traits at each elevation, then calculated multivariate plasticity across elevations as per Noble et al. (2019) using

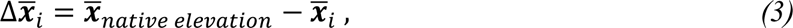

where for each species, 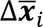 is a vector of differences in mean trait values (for all five leaf traits) between the native elevation (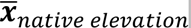) and the *i*th novel elevation (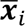).

We then tested whether plasticity in leaf traits produced phenotypes with abundant genetic variation at each elevation using the matrix projection

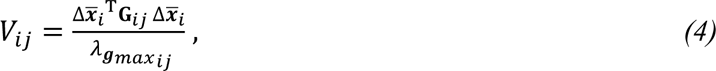

where for each species, 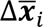 is the plasticity vector from equation 3, *T* is its transpose and **G**_*ij*_ is the *j*th MCMC sample of **G** at the *i*th novel elevation. The projection is divided by λ_**g***max ij*_ (the eigenvalue of ***g***_max_) to estimate genetic variance in the direction of leaf plasticity as a proportion of maximum available genetic variation (Noble et al. 2019). Comparing genetic variance estimated at native *versus* novel elevations tests whether changes in **G** (due to G×E) reduces or increases genetic variation in the direction of plasticity.

#### Does selection favour phenotypes with abundant genetic variation in more novel environments?

To estimate selection on leaf traits, we calculated phenotypic selection gradients (***β***) by relating traits to survival (our fitness proxy) at each elevation. However, seedlings died before leaves could be sampled from either species at 2,000m, and from *S. chrysanthemifolius* at elevations above 500m (**Fig. S2**). We were therefore limited to estimating viability selection on leaf traits of *S. aethnensis* at elevations below 2,000m, acknowledging that such selection may vary over time, and that fecundity selection also acts on leaf traits. To estimate ***β***, we applied a multiple regression to mean-standardised traits and mean-standardised (relative) survival using the package *lme4* (Bates et al. 2015) and the linear mixed model

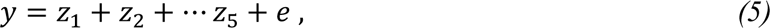

where *y* is relative survival to the end of summer, z_1-5_ are leaf traits, and *e* is the error variance. Survival was assumed to follow a normal distribution, as required by equations predicting evolutionary change from estimates of selection on continuous (polygenic) traits and genetic variation in those traits (Lande and Arnold 1983; Mitchell-Olds and Shaw 1987; Hajduk et al. 2020). We then extracted ***β*** as the vector of regression coefficients (scaled to unit length), representing the direction of viability selection on leaf traits for *S. aethnensis* at each elevation (Lande and Arnold 1983).

To test whether selection on leaf traits favours phenotypes with abundant genetic variation for *S. aethnensis* planted at novel lower elevations, we used equation 4 to project selection vectors (***β***) estimated at 500m, 1,000m and 1,500m through the corresponding G-matrices. This projection yields the proportion of genetic variation in leaf traits that lies in the direction of selection on those traits. To test whether elevational changes in **G** increased the amount of genetic variation in selection, we also projected the selection vectors through **G** estimated at the native 2,000m elevation of *S. aethnensis*. This allowed us to test how much genetic variance lay in the direction of selection at each elevation, relative to the native elevation (i.e., as if genetic variance remained unchanged across elevations). We incorporated uncertainty in estimates of selection by creating 1,000 bootstrapped samples of ***β*** using the *boot* package (Canty and Ripley 2022), and then projecting each sample through each MCMC sample of **G**.

#### How abundant is genetic variation in the directions of native phenotypes at novel elevations?

We tested whether genetic variation in leaf traits at novel elevations was abundant in the directions of native leaf phenotypes adapted to those elevations. We quantified the proportion of genetic variation in traits of *S. aethnensis* in the direction of the native phenotype of *S. chrysanthemifolius* at 500m, and genetic variation in traits of *S. chrysanthemifolius* in the direction of the native phenotype of *S. aethnensis* at 2,000m. We estimated the directions of native phenotypes using equation 3, but instead calculating differences in mean values of leaf traits between species at each elevation. We then used equation 4 to project this vector through **G** for *S. aethnensis* at 500m, and for *S. chrysanthemifolius* at 2,000m, quantifying the proportion of genetic variation in leaf traits in the directions of native phenotypes at novel elevations.

## Results

### Species differ in plastic responses of leaf traits to elevation

Species differed significantly in the multivariate plasticity of leaf traits across elevations (**Fig 3**, species×elevation Wilks’ λ=0.79, F_3,3011_=49.8, P<0.0001). At 2,000m, plasticity moved the mean leaf phenotype of *S. chrysanthemifolius* towards the native phenotype of *S. aethnensis*. At 500m, however, plasticity moved the mean phenotype of *S. aethnensis* away from the native phenotype of *S. chrysanthemifolius* (**Fig. 3**). This suggests that leaf plasticity is adaptive for *S. chrysanthemifolius* at 2,000m, but nonadaptive for *S. aethnensis* at 500m. Trait means are presented in **Fig. S4**.

**Fig. 3.**
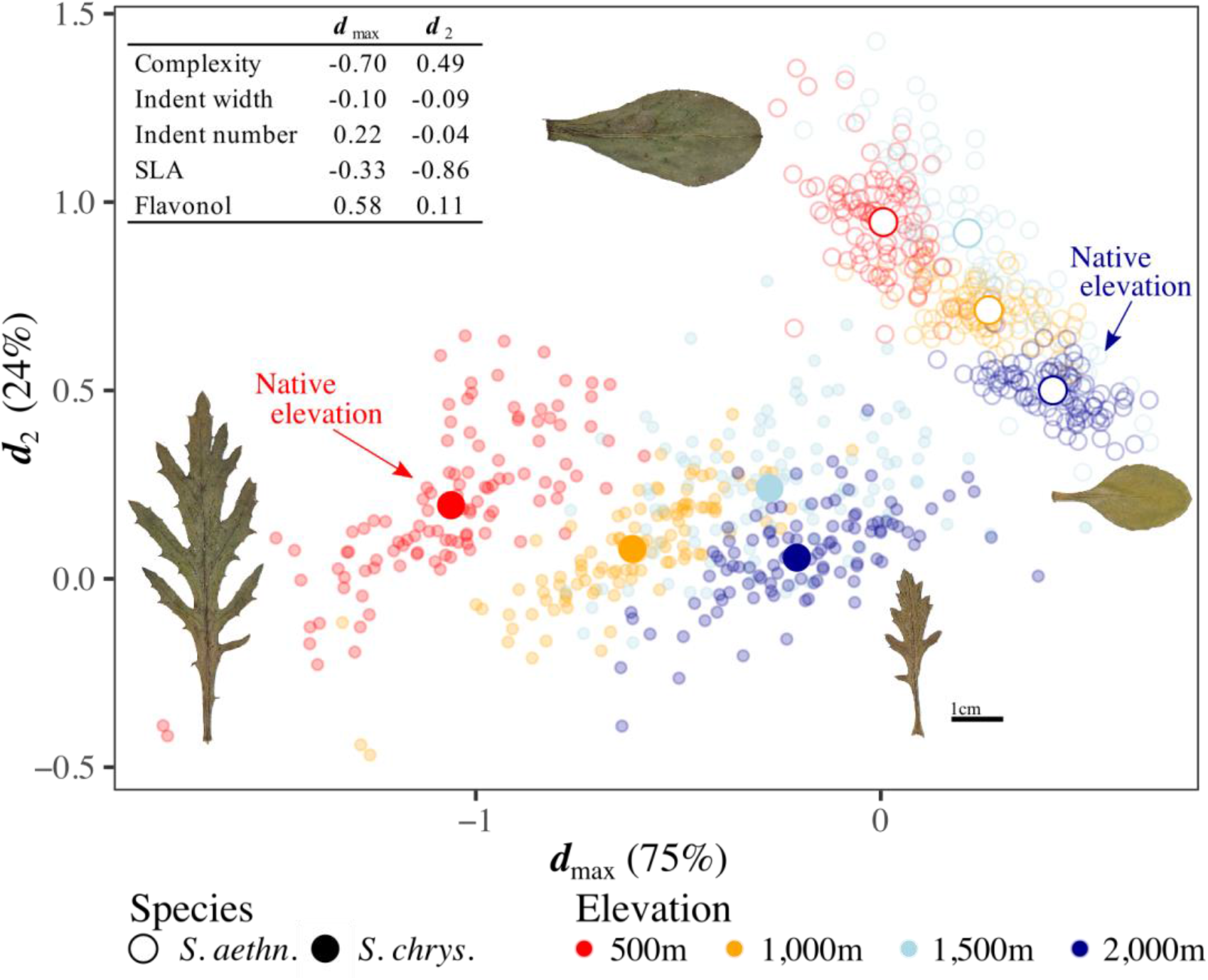
Leaf phenotypes expressed by each species at each elevation. Large, coloured circles are species’ multivariate means at each elevation (circles exceed one standard error of the mean), and leaf images show the corresponding mean phenotype at the elevational extremes for both species. Changes in mean phenotype across elevations are due to plasticity, which differs between species. Small circles are full-sibling family means (note that they do not accurately describe genetic variation in phenotypes). Mean phenotypes are calculated using the **D** matrix (see details in text), with the first two axes of **D** (***d*_max_** and ***d*_2_**, inset) summarising 99% of all differences in mean phenotypes across elevations and species. Trait loadings on each axis show how each leaf trait contributes to those differences.

### Genetic variation in leaf traits changes more across elevations than between species

For both species, estimates of additive genetic variance in leaf traits were significant at each elevation (except for SLA in *S. chrysanthemifolius* at high elevations; **Fig. S5**), and tended to be lower at 2,000m compared to other elevations (**Fig. S6**). Changes in **G** across elevations are summarised in **Fig. 4**. Total genetic variance was reduced at higher elevations for both species, and covariances weakened more at higher elevations for *S. aethnensis* compared to *S. chrysanthemifolius.* G-matrices are presented in **Table S3**.

**Fig. 4.**
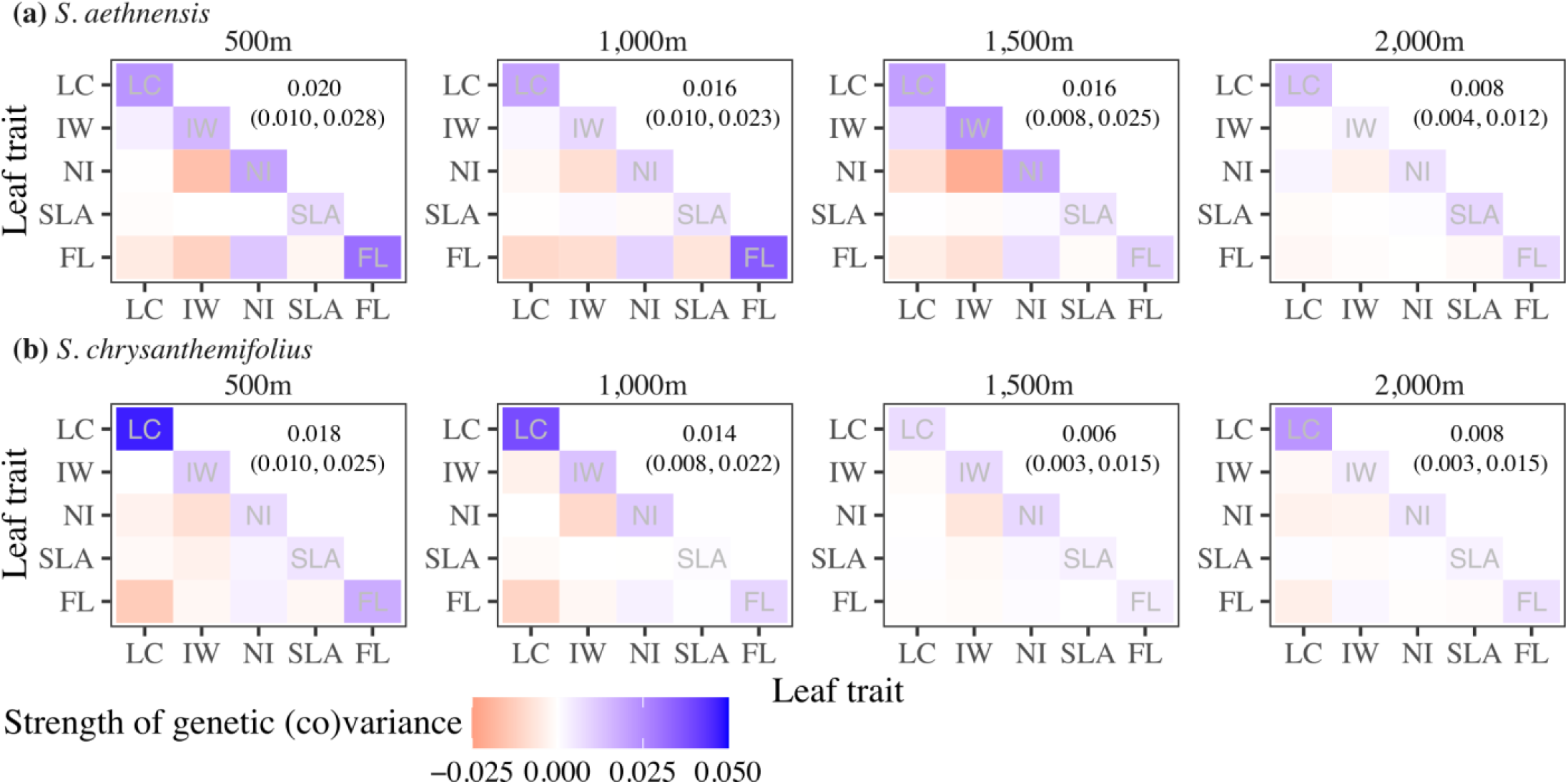
Changes in genetic variation in leaf traits of **(a)** *S. aethnensis* and **(b)** *S. chrysanthemifolius* across elevations, visualised as heat maps with elevation increasing left to right. Genetic variances of traits are on the diagonals, with the total genetic variance in each matrix presented above the diagonal (95% highest posterior density intervals in parentheses). Genetic covariances between pairs of traits are below the diagonals. Darker colours indicate stronger variances or covariances, with negative values in red and positive values in blue. Overall, genetic variation in leaf traits differs most between 2,000m and all lower elevations for *S. aethnensis*, and between 1,500m and all other elevations for *S. chrysanthemifolius*. LC=Leaf complexity, IW=Indent width, NI=Number of indents, SLA=Specific leaf area and FL=Flavonol content.

The first two eigenvectors of **G** described 62-91% of total genetic variance (**Table 1**), more than expected under the null distribution (**Fig. S7**), providing evidence that our axes of **G** are statistically significant. At most elevations, ***g***_max_ (the primary axis of **G**) described >50% of total genetic variation in leaf traits of each species, suggesting strong genetic correlations among leaf traits (**Table 1**). At 2,000m, however, ***g***_max_ described 58% genetic variation in leaf traits of *S. chrysanthemifolius*, compared to 37% for *S. aethnensis,* suggesting weaker genetic correlations for *S. aethnensis* at its native elevation.

**Table 1.** The first two eigenvectors of **G** (***g*_max_** and ***g*_2_**) for *S. aethnensis* (high elevation species) and *S. chrysanthemifolius* (low elevation species) at each transplant elevation. Eigenvectors represent directions in which leaf traits vary genetically. Trait loadings >0.2 (in bold) are interpreted as contributing substantially to each eigenvector. Eigenvalues and their 95% highest posterior density (HPD) intervals estimate additive genetic variation in these directions, which summarise 62-91% of all variation in **G** in each case.

|  | 500m |  | 1,000m |  | 1,500m |  | 2,000m |  |
| --- | --- | --- | --- | --- | --- | --- | --- | --- |
| | $g_{\max}$ | $g_2$ | $g_{\max}$ | $g_2$ | $g_{\max}$ | $g_2$ | $g_{\max}$ | $g_2$ |
| *S. aethnensis* |  |  |  |  |  |  |  |  |
| **Eigenvalue** | 0.051 | 0.023 | 0.047 | 0.017 | 0.051 | 0.016 | 0.015 | 0.010 |
| **HPD** | 0.029, 0.090 | 0.013, 0.038 | 0.026, 0.082 | 0.009, 0.029 | 0.025, 0.092 | 0.007, 0.031 | 0.009, 0.032 | 0.005, 0.018 |
| **Proportion (%)** | 52 | 23 | 59 | 22 | 63 | 20 | 37 | 25 |
| **Trait loading** |  |  |  |  |  |  |  |  |
| Complexity | -0.19 | **-0.93** | **-0.35** | **-0.89** | **-0.37** | **-0.92** | **-0.92** | 0.11 |
| Indent width | **-0.48** | 0.14 | **-0.27** | **0.27** | **-0.65** | **0.33** | 0.10 | -0.04 |
| Indent no. | **0.52** | **-0.33** | **0.29** | **-0.30** | **0.61** | **-0.21** | **-0.28** | -0.05 |
| SLA | -0.03 | 0.03 | -0.15 | 0.17 | 0.02 | -0.06 | 0.07 | **-0.80** |
| Flavonol | **0.67** | 0.09 | **0.84** | -0.14 | **0.27** | 0.04 | **0.25** | **0.58** |
| *S. chrysanthemifolius* |  |  |  |  |  |  |  |  |
| **Eigenvalue** | 0.053 | 0.019 | 0.043 | 0.023 | 0.016 | 0.008 | 0.024 | 0.009 |
| **HPD** | 0.026, 0.091 | 0.011, 0.032 | 0.021, 0.078 | 0.01, 0.034 | 0.005, 0.039 | 0.001, 0.015 | 0.008, 0.058 | 0.001, 0.018 |
| **Proportion (%)** | 59 | 22 | 59 | 32 | 50 | 25 | 58 | 22 |
| **Trait loading** |  |  |  |  |  |  |  |  |
| Complexity | **0.92** | 0.18 | **0.95** | 0.00 | -0.08 | **0.97** | **0.96** | 0.07 |
| Indent width | 0.04 | **-0.73** | -0.10 | **-0.71** | **0.69** | -0.06 | -0.08 | **0.57** |
| Indent no. | -0.10 | **0.57** | 0.01 | **0.66** | **-0.69** | -0.16 | -0.15 | **-0.64** |
| SLA | -0.02 | **0.24** | -0.03 | 0.00 | -0.17 | 0.05 | 0.04 | -0.18 |
| Flavonol | **-0.36** | **0.21** | **-0.30** | **0.24** | -0.11 | -0.18 | **-0.23** | **0.47** |

For *S. aethnensis, **g***_max_ represented a different combination of leaf traits at its native elevation (2,000m) than at lower elevations (**Table 1**). Large angles between ***g***_max_ at the native elevation compared to other elevations (θ for 2,000m-1,500m = 77.68° [39.95, 89.97 HPD]; 2,000m-1,000m = 80.91° [34.81, 89.93 HPD]; 2,000m-500m = 83.91° [36.99, 89.96 HPD]) indicated large changes in the orientation of genetic variation in leaf traits across elevations. For *S. chrysanthemifolius*, by comparison*, **g***_max_ represented a similar combination of leaf traits at all elevations except 1,500m (**Table 1**). This corresponded to small angles between ***g***_max_ at the native elevation (500m) compared to 1,000m (θ = 11.89° [4.16, 56.16 HPD]) and 2,000m (θ = 22.30° [7.94, 66.33 HPD]), but a larger angle between ***g***_max_ at the native elevation compared to 1,500m (θ = 84.24° [35.49, 90.00 HPD]).

Three (of seven) eigentensors captured significant differences in **G** across species and elevations (**Fig. S8** and **Table S4**). The first eigentensor (E1, 37.6% of all differences in **G**) described large differences in genetic variation in leaf traits across elevations, but only small differences in genetic variation between species (**Fig. 5**). By contrast, the second eigentensor (E2, 28.1% of all differences in **G**) described large differences in genetic variation in leaf traits between species, but smaller differences across elevations (**Fig. 5**). Hence, genetic variation in leaf traits differed more in response to elevation than to adaptive divergence between the two species. The third eigentensor (E3, 16% of all differences in **G**) described small differences between the two intermediate elevations (not shown).

**Fig. 5.**
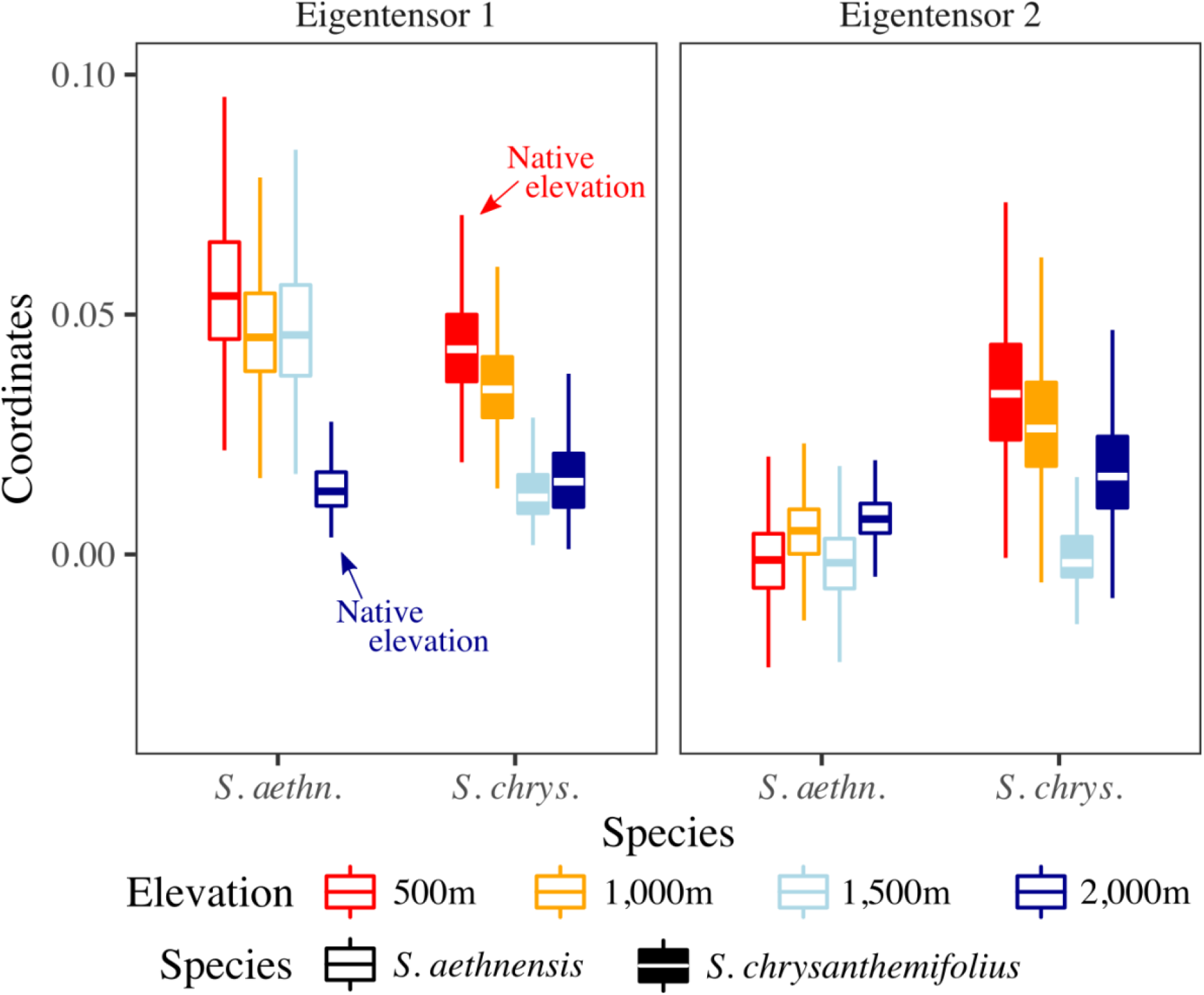
Genetic variation in leaf traits differs more across elevations than between species. Eigentensors summarise differences in **G** across elevations and between species, while coordinates indicate how much each original matrix contributes to the differences described by each eigentensor. Boxplots are posterior distributions for the matrix coordinates. Eigentensor 1 describes large elevational differences, but relatively small species differences, in genetic variation in leaf traits. Eigentensor 2 describes large species differences in genetic variation in leaf traits, which is higher at the lower elevations.

### Genetic variation in leaf traits is moderate in the direction of plasticity

For *S. aethnensis*, c.50% of the maximum amount of genetic variation available at each elevation was in the direction of leaf plasticity across elevations, except for genetic variance estimated at 1,500m where only c.10% was in the direction of plasticity (**Fig. 6a**). For *S. chrysanthemifolius*, plasticity was in a direction of the leaf phenotype containing c.50-70% of the maximum genetic variance available, and at all elevations plasticity was associated with greater genetic variance at the native elevation compared to the other elevations (**Fig. 6b**). Leaf plasticity in both species therefore occurred in a direction of the phenotype containing a moderate amount of genetic variance.

**Fig. 6.**
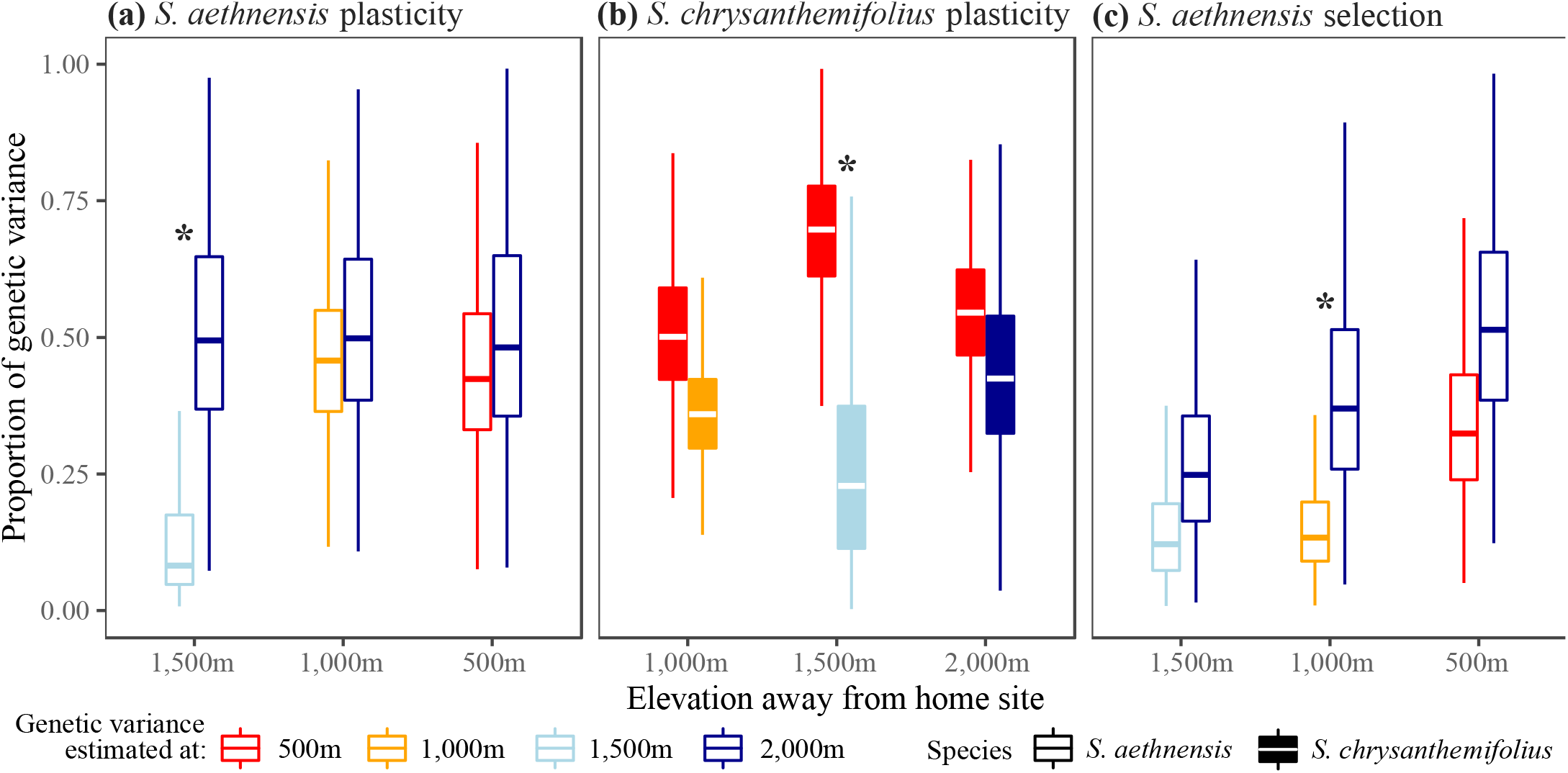
Boxplots of posterior distributions for the proportions of genetic variation in leaf traits that lie in the directions of **(a-b)** elevational plasticity in those traits and **(c)** viability selection on those traits in *S. aethnensis*. If plasticity and selection are in directions with higher genetic variation at the native elevation than at other elevations, then G×E reduces genetic variation in those directions at other elevations. For both *S. aethnensis* **(a)** and *S. chrysanthemifolius* **(b)**, plasticity changed the phenotype in a direction containing more genetic variation at the native elevation compared to more novel elevations. **(c)** For *S. aethnensis,* only a small proportion (<25%) of genetic variance lay in the direction of selection. Asterisks denote significant differences between genetic variation at the native elevation *versus* other elevations (distributions do not overlap at >90% HPD).

### Elevational changes in genetic variation in leaf traits reduces genetic variation in the direction of selection

Viability selection on leaf traits in *S. aethnensis* favoured greater SLA (*χ^2^*(1)=1.28, P<0.01) and flavonol content (*χ^2^*(1)=2.00, P<0.01) at 1,500m, greater leaf complexity (*χ^2^*(1)=1.71, P<0.01) and SLA (*χ^2^*(1)=3.00, P<0.01) at 1,000m, and greater leaf complexity (*χ^2^*(1)=132.46, P<0.01) and flavonol content (*χ^2^*(1)=41.50, P<0.01) at 500m (**Table S5** and **Fig. S9**).

For *S. aethnensis* planted at lower elevations (500-1,500m), a small proportion (<25%) of genetic variation in leaf traits lay in the direction of viability selection (***β***) (**Fig. 6c**). Furthermore, the direction of viability selection at each elevation described more genetic variation at the native elevation, suggesting that changes in genetic variance across elevations reduced the amount of genetic variation in the direction of selection (**Fig. 6c**). Analysing genotypic selection gradients, estimated using family means for traits and survival, gave similar results (**Fig. S10**).

### Genetic variation in leaf traits is abundant in the direction of the native phenotype of only one species

For *S. aethnensis* planted at the novel 500m elevation, c.40% of genetic variation in leaf traits was in the direction of the native phenotype of *S. chrysanthemifolius.* Furthermore, compared to genetic variance estimated at 500m, a greater proportion of genetic variance estimated at the native 2,000m elevation (c.60%) was in the direction of the native phenotype at 500m. Hence, when *S. aethnensis* was planted at the novel elevation, G×E in leaf traits created changes in **G** from the native 2,000m to the novel 500m that reduced genetic variation in the direction of the native phenotype adapted to 500m (**Fig. 7**). By contrast, for *S. chrysanthemifolius* at 2,000m, a greater proportion (c.70%) of genetic variance was in the direction of the native phenotype of *S. aethnensis* (**Fig. 7**), and this was similar for genetic variance estimated at 500m and 2,000m. Therefore, although **G** changed less across elevations for *S. chrysanthemifolius*, more genetic variance was in the direction of the native phenotype at the novel elevation.

**Fig. 7.**
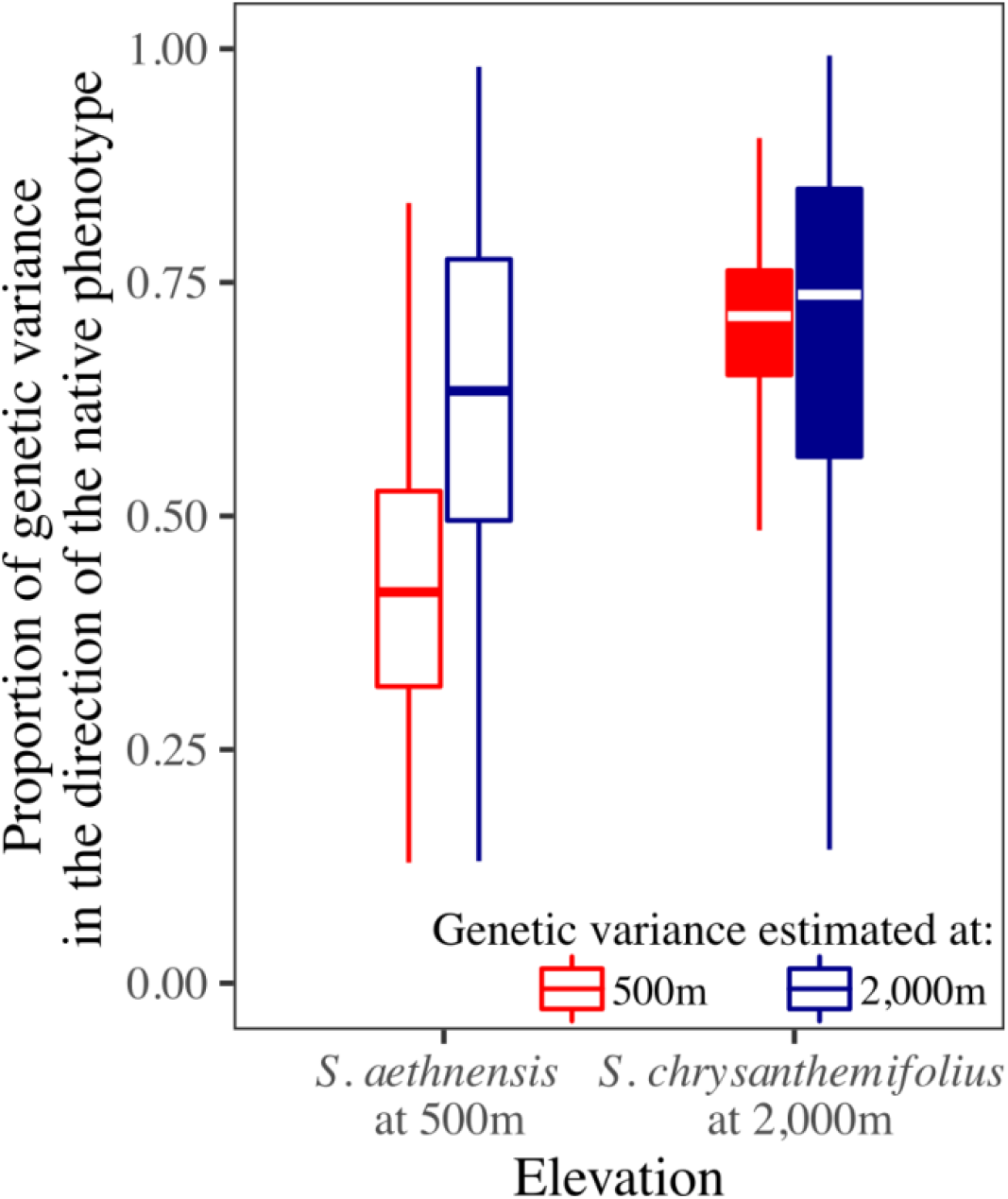
Boxplots of posterior distributions for the proportions of genetic variation in leaf traits that lie in the direction of the native phenotype of each species planted at both native elevations. At 500m, genetic variation in leaf traits of *S. aethnensis* aligns more poorly with the native phenotype of *S. chrysanthemifolius*, when compared to genetic variance estimated at the native 2,000m elevation. At 2000m, genetic variation in leaf traits of *S. chrysanthemifolius* (estimated at both the native and novel elevations) align more closely with the native phenotype of *S. aethnensis*.

## Discussion

Field studies combining estimates of plasticity, selection, and genetic variation are exceedingly rare. We used a breeding design to produce seeds for two closely related but ecologically distinct Sicilian *Senecio* species, which we reciprocally planted across an elevational gradient. We estimated plasticity, viability selection, and genetic variation for five ecologically important leaf traits on emergent seedlings. The two species showed differences in leaf plasticity (**Fig. 3**), with *S. chrysanthemifolius* showing more evidence of adaptive plasticity (Walter et al. 2022a). Higher elevations were associated with less genetic variation in leaf phenotypes (**Fig. 4**) and meant that elevation created larger differences in genetic variation in leaf traits than differences between species (**Fig. 5**). Genetic correlations among leaf traits showed less change across elevations for *S. chrysanthemifolius* when compared to *S. aethnensis,* which meant that genetic variance in leaf phenotypes changed more across elevations for *S. aethnensis* than *S. chrysanthemifolius.* For both species, plasticity across elevations produced leaf phenotypes that contained a moderate amount (50-60%) of genetic variance (**Fig. 6**). For *S. aethnensis*, changes in genetic variance in leaf traits at lower elevations reduced the amount of genetic variance in the direction of phenotypic selection on seedling viability (<25%; **Fig. 6c**) and the native phenotype of *S. chrysanthemifolius* at 500m (c.40% at 500m; **Fig. 7**). By contrast, *S. chrysanthemifolius* showed a greater proportion of genetic variance in leaf traits in the direction of the native phenotype at the novel 2,000m elevation (c.70%; **Fig. 7**). These results suggest that for *S. chrysanthemifolius* at the novel 2,000m elevation, changes in genetic variance were smaller and associated with the direction of the native phenotype at that elevation (reflecting the scenario in **Fig. 1b**). By contrast, for *S. aethnensis* at the novel 500m elevation, larger changes in genetic variance were observed, but were associated with less genetic variance in the direction of viability selection and the native phenotype (reflecting the scenario in **Fig. 1d**). Our results therefore suggest that nonadaptive plasticity and large environmental effects on genetic variance introduce genetic constraints for adaptation in novel environments.

### Moderate amounts of genetic variance in the direction of plasticity

We found moderate amounts of genetic variance in the direction of plasticity, providing some support for a meta-analysis that suggested plasticity is biased in directions of phenotypes with large amounts of additive genetic variance (Noble et al. 2019). In this meta-analysis, however, the amount of genetic variance in the direction of plasticity varied across species, traits and environments. The alignment between plasticity and genetic variation could differ among studies because species (and populations) vary in their amount of genetic variation in plasticity. During adaptation, if the environment is predictable (as required for adaptive plasticity to evolve; Leung et al. 2020) and selection on plasticity is strong, then adaptation is likely to erode genetic variation in plasticity (Oostra et al. 2018; Hoffmann and Bridle 2022), which could result in plasticity in the direction favoured by historical selection rather than that of contemporary genetic variation. This could explain our observation of only a moderate amount of genetic variance in the direction of plasticity. Resolving how and when genetic variation aligns with plasticity (and understanding whether plasticity is adaptive) will reveal when selection on existing plastic responses could promote adaptation to a novel environment. Regardless, our results suggest that if genetic variation aligns with plasticity that is nonadaptive (as was the case for the high elevation *Senecio*), then it is likely that plasticity will create genetic constraints that prevent rapid adaptation to novel environments.

### Environmental effects on genetic variance

To better understand the potential for rapid adaptation, it is important to understand how genotypes across a species’ range vary in their responses to environmental variation, and whether populations harbour genetic variation that could help align **G** with selection in novel environments (Lande 2009; Chevin and Lande 2011; Lind et al. 2015; Bridle and Hoffmann 2022; Walter et al. 2023). In our results, G×E produced larger changes in genetic variation in leaf traits across elevations than between two closely related species of *Senecio*. In novel low elevations, G×E in the high elevation species created larger changes in genetic variation in leaf traits that reduced the amount of genetic variation in the direction of selection. By contrast, the low elevation species showed smaller changes in genetic variation at higher elevations that resulted in greater genetic variation in the direction of the native phenotype. Wood and Brodie III (2015) found evidence that **G** is likely to be affected by the environment as much as by evolution, although it was unclear why **G** changed with the environment. We help to resolve this by demonstrating that novel environments not only created larger changes in genetic variance in leaf phenotypes than evolutionary history, but also that such changes can be the result of nonadaptive genotype-by-environment interactions (G×E) in response to stress when the high elevation species experience low elevations. Changes in **G** may often be maladaptive in novel environments because plasticity has not evolved to suit those conditions. *Senecio aethnensis* shows evidence of greater genomic diversity than *S. chrysanthemifolius* (Chapman et al. 2013), suggesting that G×E underlying the large elevational changes in genetic variation in leaf traits observed could be because *S. aethnensis* originated from a larger ancestral population. Given that evolutionary history likely determines G×E, the extent to which G×E is adaptive in novel environments is likely to be species-specific.

### Estimating genetic variance and selection when there is viability selection

Although we provide substantial evidence for elevational changes in genetic variance for both species, we could not measure all individuals before high mortality occurred. Our estimates of genetic variance could therefore be biased if viability selection removed certain leaf phenotypes before they could be measured (Hadfield 2008). This issue is notoriously difficult to avoid in field experiments, but given that we measured individuals from almost all families of *S. chrysanthemifolius*, and all families of *S. aethnensis*, before substantial mortality occurred, our estimates should be robust. Further, to ensure adequate replication of families across elevations, it was necessary to plant seeds close to each other. This meant that we could only measure early stages of development because plants grew into one another as they reached maturity. Our estimates of selection are therefore limited to selection operating through early survival and exclude later episodes of selection operating during reproduction. Nevertheless, populations experiencing novel environments are likely to be under stronger viability than fecundity selection, at least during the initial colonisation of a novel environment (Hadfield 2008; Mittell and Morrissey 2023).

## Conclusion

We show that environmental effects on genetic variance in leaf traits are likely to introduce genetic constraints for adapting to novel environments. When species encounter a novel environment, the extent to which their existing plasticity is adaptive, and their potential for adaptation to the new environment, will be species-specific (Walter et al. 2022a). For both *Senecio* species studied here, plasticity produced leaf phenotypes with moderately high amounts of genetic variation, suggesting that the direction of plasticity will, to some extent, determine the direction of phenotypic evolution and will likely determine the strength of genetic constraints that will slow adaptation and make population extinction more likely than evolutionary rescue. Genetic variance in leaf traits changed more across elevations for the high elevation species (than the low elevation species), with such changes reducing the amount of genetic variation in the direction of viability selection on leaf traits. Therefore, although we previously showed that genetic variance in survival increased in both species at novel elevations, suggesting that adaptive potential increases in novel environments for both species (Walter et al. 2022b), it is likely that the high-elevation species will face genetic constraints that will make it difficult for rapid adaptation to occur at lower elevations, and by extension, warmer conditions created by climate change. By contrast, rapid adaptation to the high-elevation habitat should be more likely because genetic variation in *S. chrysanthemifolius* leaf phenotypes at 2,000m aligned more closely with plasticity and the native phenotype. Such variation in responses to warmer versus cooler environments, even between closely related species, makes predicting adaptation to novel environments challenging. Nevertheless, our results support evidence that limits to adaptation and plasticity are stronger at warmer range margins than at cooler margins (Kellermann et al. 2012; van Heerwaarden et al. 2016; Sheth and Angert 2018; Anderson and Wadgymar 2020; Arnold et al. 2022).

## Acknowledgements

We are very grateful to Piante Faro (Giarre, Italy) for providing us with glasshouse facilities. We thank Giuseppe Riggio for generously providing us access to the 1,000m field site. This work was carried out using the computational facilities of the Advanced Computing Research Centre, University of Bristol. This work was supported by joint NERC grants NE/P001793/1 and NE/P002145/1 awarded to JB and SH. GW was supported by an Australian Research Council early career (DECRA) fellowship DE200101019.

## Author contributions

GW designed the study with AC, SH, SC and JB. GW, DT, ES, MM, GP and AC conducted the experiments, collected and curated the data. GW analysed the data and wrote the paper with help from KM, JC and input from all authors.

## Data availability

Upon acceptance, all data and code will be deposited with Dryad.

## Notes

### Competing Interest Statement

The authors have declared no competing interest.

### Summary of Updates

Revised introduction, results and discussion.

