## Supplementary material for "Environmental effects on genetic variance are likely to constrain adaptation in novel environments"

**Environmental effects on genetic variance and the alignment with plasticity and selection**

**Contents**

**Table S1** Sampling locations
**Fig. S1** Map of sampling locations **Fig. S2** Demography during the field experiment **Table S2** Numbers of individuals that were measured in the field experiment
**Fig. S3** Description of the covariance tensor approach **Fig. S4** Elevational change in mean for all five leaf traits.
**Fig. S5** Significance test for estimates of genetic variance for each trait **Fig. S6** Posterior distribution of genetic variance for each trait
**Table S3** G-matrices for all elevations and for both species
**Fig. S7** Significance test for eigenvectors of **G**
**Fig. S8** Significant eigentensors
**Table S4** Summary of the covariance tensor analysis
**Table S5** Coefficients for estimates of selection
**Fig. S9** Visualising the selection gradients
**Methods S1** Visualizing differences in mean multivariate phenotype

**Table S1:** Location of sampled individuals for the parental generation of the breeding design for each species. The final two column denote the number of individuals used as sires and dams in the breeding design.

| **Species** | **Site** | **Elevation** | **Latitude** | **Longitude** | **# Sires** | **# Dams** |
| --- | --- | --- | --- | --- | --- | --- |
| *S. aethn.* | Etna South | 2,600m | 37°43'13.28"N | 15° 0'3.54"E | 1 | 1 |
|  |  | 2,500m | 37°43'3.80"N | 14°59'59.20"E | 8 | 7 |
|  |  | 2,400m | 37°42'46.50"N | 14°59'41.30"E | 5 | 5 |
|  |  | 2,200m | 37°42'24.82"N | 14°59'42.69"E | 2 | 5 |
|  | Etna North | 2,600m | 37°46'39.90"N | 15° 0'23.00"E | 8 | 4 |
|  |  | 2,500m | 37°46'53.70"N | 15° 0'28.80"E | 3 | 2 |
|  |  | 2,400m | 37°47'7.46"N | 15° 0'35.05"E | 4 | 8 |
|  |  | 2,200m | 37°47'32.82"N | 15° 1'14.53"E | 5 | 3 |
|  | **Totals** |  |  |  | **36** | **35** |
| *S. chrys.* | Bonnano | 790m | 37°38'24.92"N | 15° 2'50.80"E | 9 | 11 |
|  | Cacciola | 680m | 37°37'31.32"N | 15° 3'26.71"E | 7 | 6 |
|  | Poggofelice | 526m | 37°39'44.31"N | 15° 5'48.55"E | 6 | 6 |
|  | Spina | 730m | 37°39'19.27"N | 15° 4'30.92"E | 10 | 10 |
|  | Trecastagni | 571m | 37°36'46.67"N | 15° 4'29.64"E | 6 | 5 |
|  | **Totals** |  |  |  | **38** | **38** |

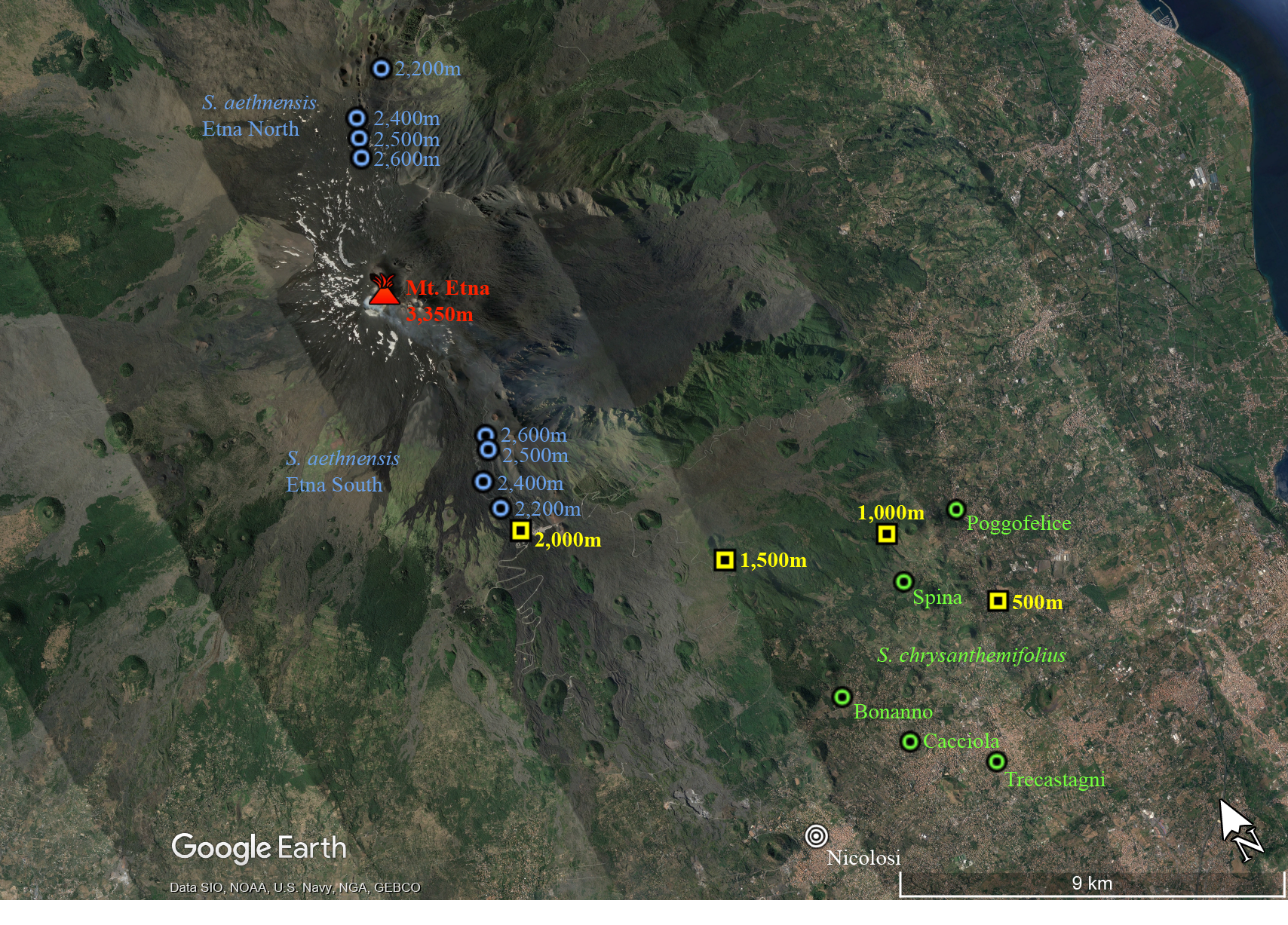

**Fig. S1** Map with the locations of transplant sites (yellow), and the sites that the parental genotypes were sampled from for both *S. aethnensis* (blue) and *S. chrysanthemifolius* (green).

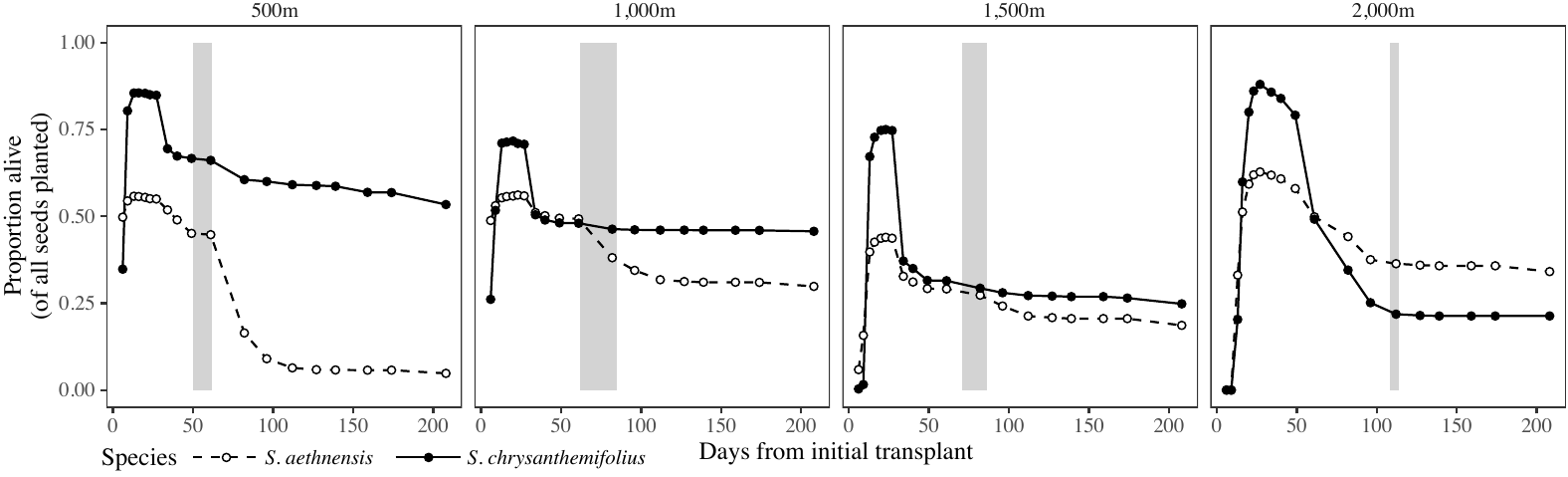

**Fig. S2** Proportion of plants alive at each census day, for each transplant elevation. Filled circles and solid lines represent *S. chrysanthemifolius*, while unfilled circles and broken lines represent *S. aethnensis*. Grey bars denote the time period during which leaf measurements were taken for each elevation.

**Table S2:** The number of seedlings measured for leaf traits at each elevation and for each species.

| **Species** | **Elevation** | **Number of seedlings emerged** | **Number of seedlings measured** | **Proportion of plants that were measured** | **Average number of individuals measured per family (± 1 SD)** |
| --- | --- | --- | --- | --- | --- |
| *S. aethn.* | 500m | 1224 | 689 | 0.56 | 7.33 (3.9) |
|  | 1,000m | 1215 | 786 | 0.65 | 8.36 (3.8) |
|  | 1,500m | 990 | 442 | 0.45 | 4.70 (3.1) |
|  | 2,000m | 1393 | 644 | 0.46 | 6.85 (3.5) |
|  | **Totals** | **4822** | **2561** | **0.53** |  |
| *S. chrys.* | 500m | 2331 | 1683 | 0.72 | 15.58 (3.4) |
|  | 1,000m | 1912 | 1144 | 0.60 | 10.59 (3.4) |
|  | 1,500m | 2061 | 589 | 0.29 | 5.45 (2.3) |
|  | 2,000m | 2414 | 482 | 0.20 | 4.46 (2.2) |
|  | **Totals** | **8718** | **3898** | **0.45** |  |

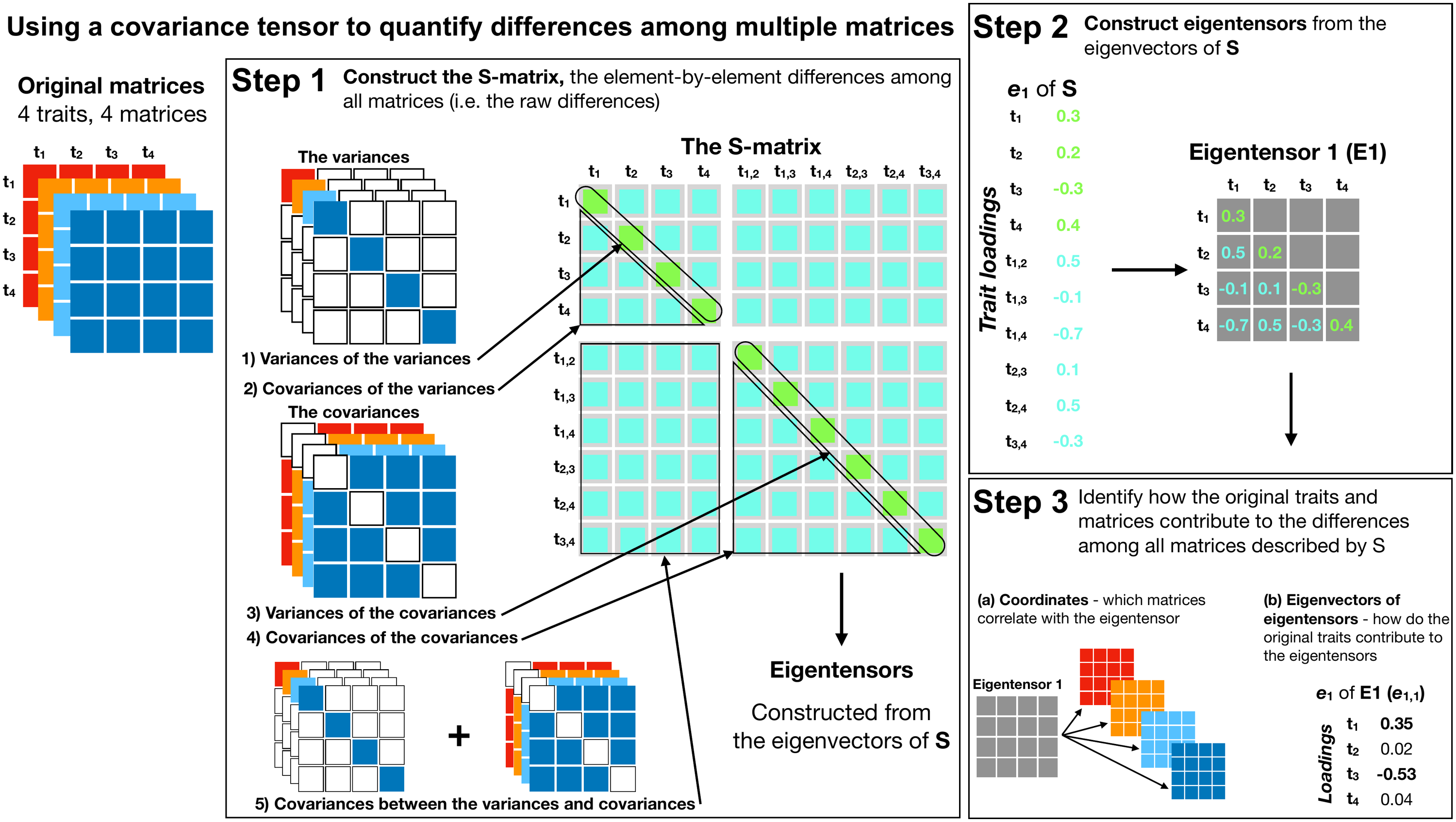

**Fig. S3** Summary of the covariance tensor approach for comparing multiple matrices.

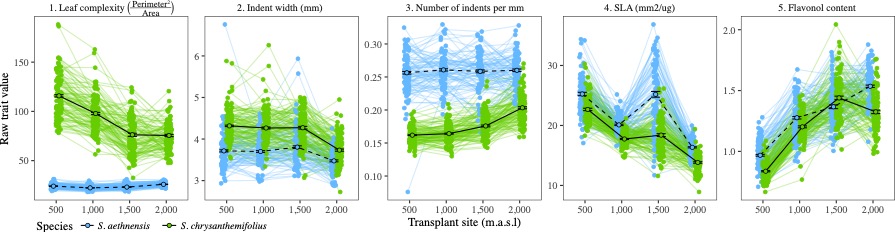

**Fig. S4** Change in univariate traits across elevation for both species. *Senecio aethnensis* is represented in blue, and *S. chrysanthemifolius* in green. Black circles and lines represent the overall mean for each species at each elevation (confidence intervals represent one standard error). Coloured lines represent the mean for each full-sibling family at each elevation.
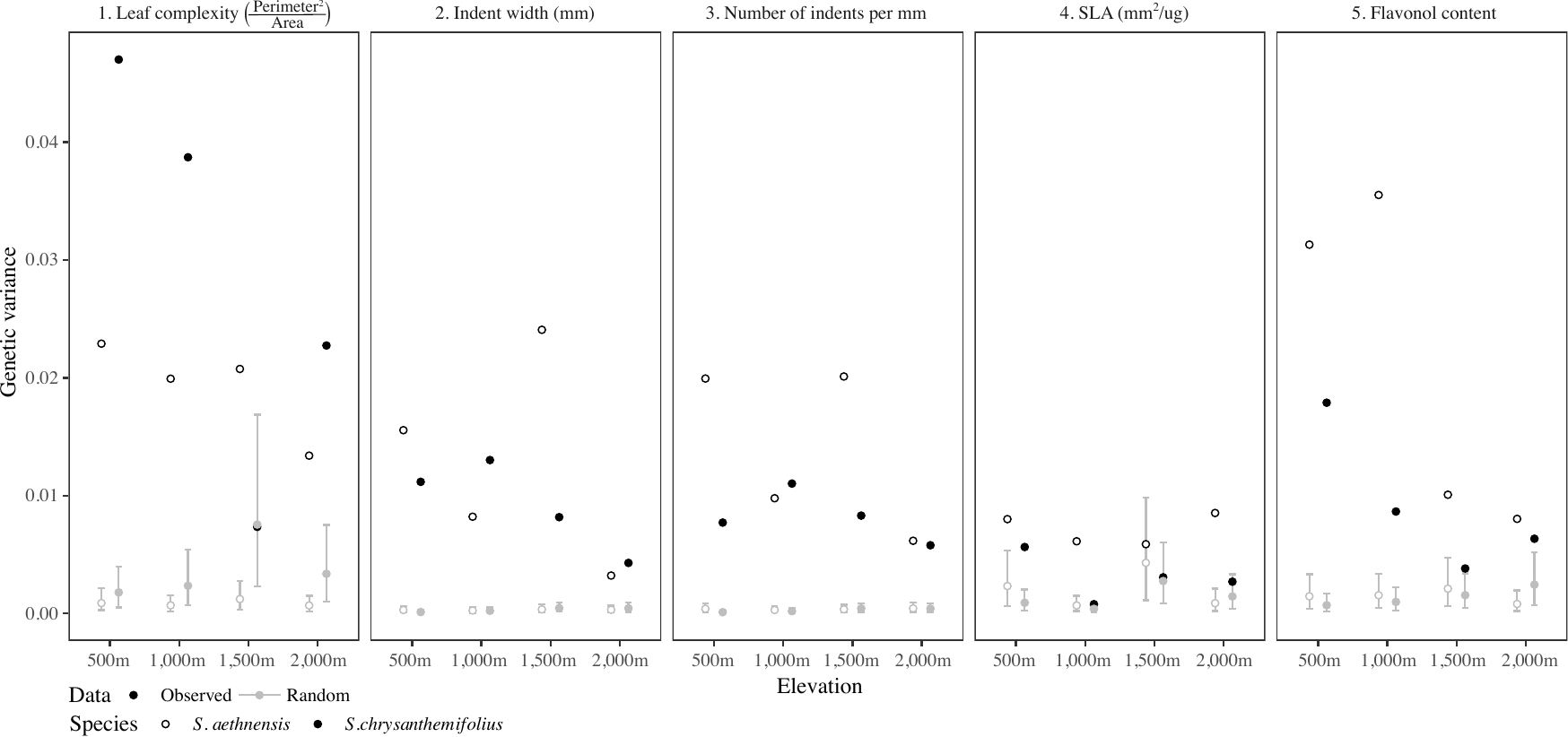

**Fig. S5** Observed estimates of genetic variance (black circles) are greater than the null distribution (grey circles and 95% HPD interval) for most traits. Open circles represent *S. aethensis,* and closed circles *S. chrysanthemifolius.* Observed estimates represent the posterior mean of the observed models, while the random distribution is the distribution of the 1,000 models, each conducted on a randomisation of the pedigree and taking the mean from each model. Leaf complexity at 1,500m and SLA estimated at higher elevations for *S. chrysanthemifolius* were the only traits that showed less observed genetic variance than expected under random sampling*.*

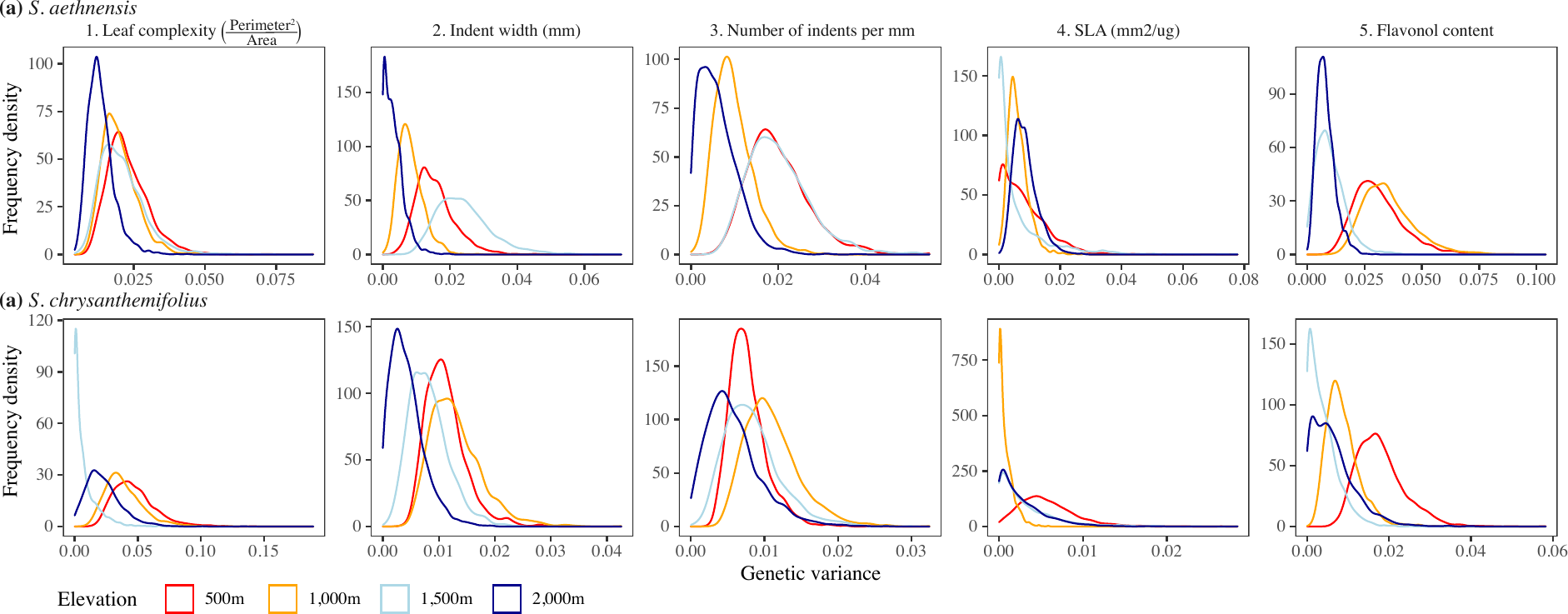

**Fig. S6** Posterior distributions for estimates of genetic variance for each trait and species. Colours represent the different elevations (red=500m, orange=1,000m, light blue=1,500m and dark blue=2,000m). While the estimates of genetic variance for some traits are bounded by zero, genetic variance for most traits are normally distributed. **Table S3** G-matrices estimated at each elevation for (**a)** *S. aethnensis* and (**b)** *S. chrysanthemifolius*. Grey shading denotes the genetic variances along the diagonal. Genetic covariances are presented above the diagonal, and genetic correlations below the diagonal. Numbers in parentheses represent the 95% HPD interval for each parameter estimated. Numbers in bold denote genetic correlations with an absolute magnitude greater than 0.2 to aide interpretation. Traits: LC=leaf complexity, NI=number of leaf indents, IW=indent width, SLA=specific leaf area, and FL=flavonol content.

| **(a) *S. aethnensis*** | | |  | |  | |  | |
| --- | --- | --- | --- | --- | --- | --- | --- | --- |
|  | **500m** | | | | | | | |
|  | LC | IW | | NI | | SLA | | FL |
| LC | 0.023 (0.011, 0.033) | 0.003 (-0.004, 0.011) | | 0.000 (-0.008, 0.007) | | -0.001 (-0.006, 0.005) | | -0.005 (-0.015, 0.005) |
| IW | 0.18 (-0.16, 0.53) | 0.016 (0.007, 0.024) | | -0.016 (-0.025, -0.007) | | 0.000 (-0.006, 0.006) | | -0.012 (-0.022, -0.003) |
| NI | 0.00 (-0.36, 0.36) | **-0.92 (-0.98, -0.87)** | | 0.02 (0.008, 0.03) | | 0.000 (-0.006, 0.007) | | 0.012 (0.003, 0.023) |
| SLA | -0.06 (-0.53, 0.42) | 0.01 (-0.68, 0.77) | | 0.00 (-0.75, 0.67) | | 0.008 (0, 0.018) | | -0.002 (-0.011, 0.005) |
| FL | **-0.20 (-0.52, 0.14)** | **-0.54 (-0.86, -0.25)** | | **0.50 (0.19, 0.81)** | | -0.14 (-0.72, 0.38) | | 0.031 (0.014, 0.047) |
|  | **1,000m** | | | | | | | |
| LC | 0.02 (0.01, 0.029) | 0.002 (-0.003, 0.007) | | -0.002 (-0.007, 0.004) | | 0.000 (-0.005, 0.004) | | -0.010 (-0.019, -0.001) |
| IW | 0.13 (-0.22, 0.51) | 0.008 (0.003, 0.014) | | -0.008 (-0.014, -0.002) | | 0.001 (-0.002, 0.004) | | -0.009 (-0.016, -0.001) |
| NI | -0.12 (-0.49, 0.23) | **-0.94 (-1, -0.88)** | | 0.01 (0.003, 0.016) | | -0.001 (-0.005, 0.003) | | 0.009 (0.001, 0.017) |
| SLA | -0.02 (-0.38, 0.37) | 0.18 (-0.33, 0.67) | | -0.17 (-0.72, 0.29) | | 0.006 (0.001, 0.011) | | -0.007 (-0.014, -0.001) |
| FL | **-0.38 (-0.69, -0.11)** | **-0.51 (-0.83, -0.18)** | | **0.51 (0.16, 0.81)** | | **-0.47 (-0.78, -0.14)** | | 0.036 (0.018, 0.052) |
|  | **1,500m** | | | | | | | |
| LC | 0.021 (0.009, 0.031) | 0.008 (-0.001, 0.016) | | -0.008 (-0.017, 0) | | 0.000 (-0.006, 0.006) | | -0.005 (-0.011, 0.002) |
| IW | **0.33 (-0.02, 0.65)** | 0.024 (0.012, 0.036) | | -0.021 (-0.032, -0.01) | | -0.001 (-0.011, 0.008) | | -0.008 (-0.017, 0.001) |
| NI | **-0.40 (-0.73, -0.09)** | **-0.97 (-1, -0.94)** | | 0.02 (0.008, 0.03) | | 0.001 (-0.006, 0.011) | | 0.007 (-0.001, 0.016) |
| SLA | 0.01 (-0.64, 0.63) | -0.09 (-0.99, 0.83) | | 0.10 (-0.84, 0.99) | | 0.006 (0, 0.017) | | -0.001 (-0.006, 0.004) |
| FL | **-0.33 (-0.78, 0.05)** | **-0.54 (-0.99, -0.08)** | | **0.54 (0.07, 1)** | | -0.06 (-0.8, 0.76) | | 0.01 (0, 0.018) |
|  | **2,000m** | | | | | | | |
| LC | 0.013 (0.006, 0.02) | 0.000 (-0.003, 0.003) | | 0.002 (-0.002, 0.006) | | -0.001 (-0.005, 0.003) | | -0.002 (-0.007, 0.002) |
| IW | -0.07 (-0.58, 0.45) | 0.003 (0, 0.007) | | -0.004 (-0.008, 0) | | 0.000 (-0.003, 0.003) | | -0.001 (-0.004, 0.002) |
| NI | **0.28 (-0.11, 0.74)** | **-0.67 (-0.98, -0.18)** | | 0.006 (0, 0.012) | | 0.001 (-0.003, 0.004) | | 0.000 (-0.004, 0.004) |
| SLA | -0.11 (-0.52, 0.25) | -0.05 (-0.58, 0.51) | | 0.11 (-0.38, 0.63) | | 0.009 (0.003, 0.015) | | -0.002 (-0.005, 0.002) |
| FL | -0.19 (-0.54, 0.19) | -0.14 (-0.73, 0.44) | | -0.06 (-0.61, 0.49) | | -0.18 (-0.62, 0.24) | | 0.008 (0.002, 0.014) |

**Table S3 (*continued*)**

| **(b) *S. chrysanthemifolius*** | | |  | |  | |  | |
| --- | --- | --- | --- | --- | --- | --- | --- | --- |
|  | **500m** | | | | | | | |
|  | LC | IW | | NI | | SLA | | FL |
| LC | 0.047 (0.023, 0.072) | 0.000 (-0.009, 0.008) | | -0.003 (-0.01, 0.004) | | -0.002 (-0.009, 0.005) | | -0.013 (-0.023, -0.002) |
| IW | 0.01 (-0.35, 0.37) | 0.011 (0.006, 0.016) | | -0.008 (-0.012, -0.004) | | -0.004 (-0.007, 0) | | -0.002 (-0.007, 0.003) |
| NI | -0.17 (-0.54, 0.16) | **-0.87 (-0.97, -0.79)** | | 0.008 (0.004, 0.011) | | 0.002 (-0.001, 0.005) | | 0.003 (-0.001, 0.008) |
| SLA | -0.08 (-0.53, 0.32) | **-0.49 (-0.88, -0.12)** | | **0.37 (-0.04, 0.8)** | | 0.006 (0, 0.01) | | -0.002 (-0.007, 0.002) |
| FL | **-0.47 (-0.75, -0.18)** | -0.15 (-0.48, 0.18) | | **0.26 (-0.06, 0.58)** | | **-0.21 (-0.62, 0.21)** | | 0.018 (0.009, 0.027) |
|  | **1,000m** | | | | | | | |
| LC | 0.039 (0.016, 0.061) | -0.004 (-0.013, 0.006) | | 0.000 (-0.008, 0.008) | | -0.001 (-0.005, 0.002) | | -0.011 (-0.019, -0.003) |
| IW | -0.16 (-0.56, 0.21) | 0.013 (0.006, 0.019) | | -0.01 (-0.015, -0.004) | | 0.000 (-0.002, 0.002) | | -0.002 (-0.007, 0.002) |
| NI | 0.02 (-0.33, 0.45) | **-0.83 (-0.95, -0.71)** | | 0.011 (0.005, 0.016) | | 0.000 (-0.002, 0.002) | | 0.003 (-0.001, 0.007) |
| SLA | **-0.24 (-0.87, 0.32)** | 0.00 (-0.66, 0.77) | | -0.06 (-0.86, 0.57) | | 0.001 (0, 0.002) | | 0.000 (-0.001, 0.002) |
| FL | **-0.61 (-0.89, -0.32)** | **-0.21 (-0.61, 0.2)** | | **0.29 (-0.07, 0.69)** | | 0.13 (-0.46, 0.82) | | 0.009 (0.002, 0.014) |
|  | **1,500m** | | | | | | | |
| LC | 0.007 (0, 0.02) | -0.001 (-0.008, 0.004) | | 0.000 (-0.006, 0.006) | | 0.000 (-0.003, 0.004) | | -0.001 (-0.004, 0.003) |
| IW | -0.13 (-0.95, 0.67) | 0.008 (0.003, 0.014) | | -0.007 (-0.012, -0.002) | | -0.002 (-0.005, 0.002) | | -0.001 (-0.004, 0.002) |
| NI | 0.01 (-0.75, 0.9) | **-0.79 (-0.96, -0.64)** | | 0.008 (0.003, 0.014) | | 0.001 (-0.002, 0.005) | | 0.001 (-0.003, 0.004) |
| SLA | 0.06 (-0.74, 0.79) | **-0.35 (-0.98, 0.38)** | | **0.29 (-0.48, 0.92)** | | 0.003 (0, 0.007) | | 0.000 (-0.002, 0.002) |
| FL | -0.08 (-0.82, 0.6) | **-0.23 (-0.9, 0.43)** | | 0.18 (-0.42, 0.91) | | 0.05 (-0.64, 0.76) | | 0.004 (0, 0.009) |
|  | **2,000m** | | | | | | | |
| LC | 0.023 (0, 0.041) | -0.002 (-0.008, 0.004) | | -0.003 (-0.011, 0.004) | | 0.001 (-0.004, 0.006) | | -0.004 (-0.012, 0.003) |
| IW | **-0.22 (-0.86, 0.38)** | 0.004 (0, 0.008) | | -0.003 (-0.007, 0.001) | | -0.001 (-0.003, 0.001) | | 0.002 (-0.001, 0.005) |
| NI | **-0.22 (-0.8, 0.36)** | **-0.49 (-0.92, 0.03)** | | 0.006 (0, 0.011) | | 0.000 (-0.002, 0.003) | | -0.001 (-0.004, 0.003) |
| SLA | 0.12 (-0.51, 0.81) | **-0.30 (-0.95, 0.31)** | | 0.08 (-0.59, 0.79) | | 0.003 (0, 0.007) | | -0.001 (-0.004, 0.001) |
| FL | **-0.33 (-0.86, 0.23)** | **0.39 (-0.18, 0.95)** | | -0.12 (-0.8, 0.49) | | **-0.23 (-0.85, 0.43)** | | 0.006 (0, 0.013) |

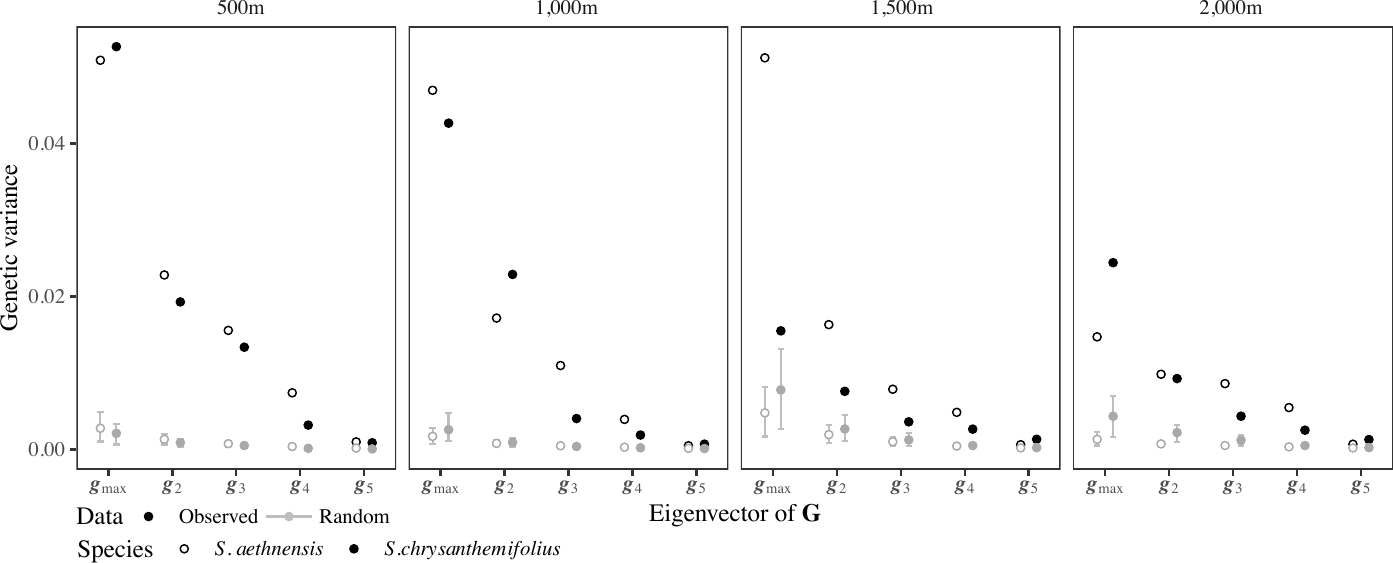

**Fig. S7** The first three eigenvectors of observed **G** (black circles) that describe >90% of total genetic variance also captured more genetic variance than expected under random sampling (grey circles, credible intervals and grey shading). Open circles represent *S. aethnensis,* and closed circles represent *S. chrysanthemifolius*. Credible intervals represent the 95% HPD (Highest Posterior Density) intervals for the models applied to the 1,000 randomisations of the data.

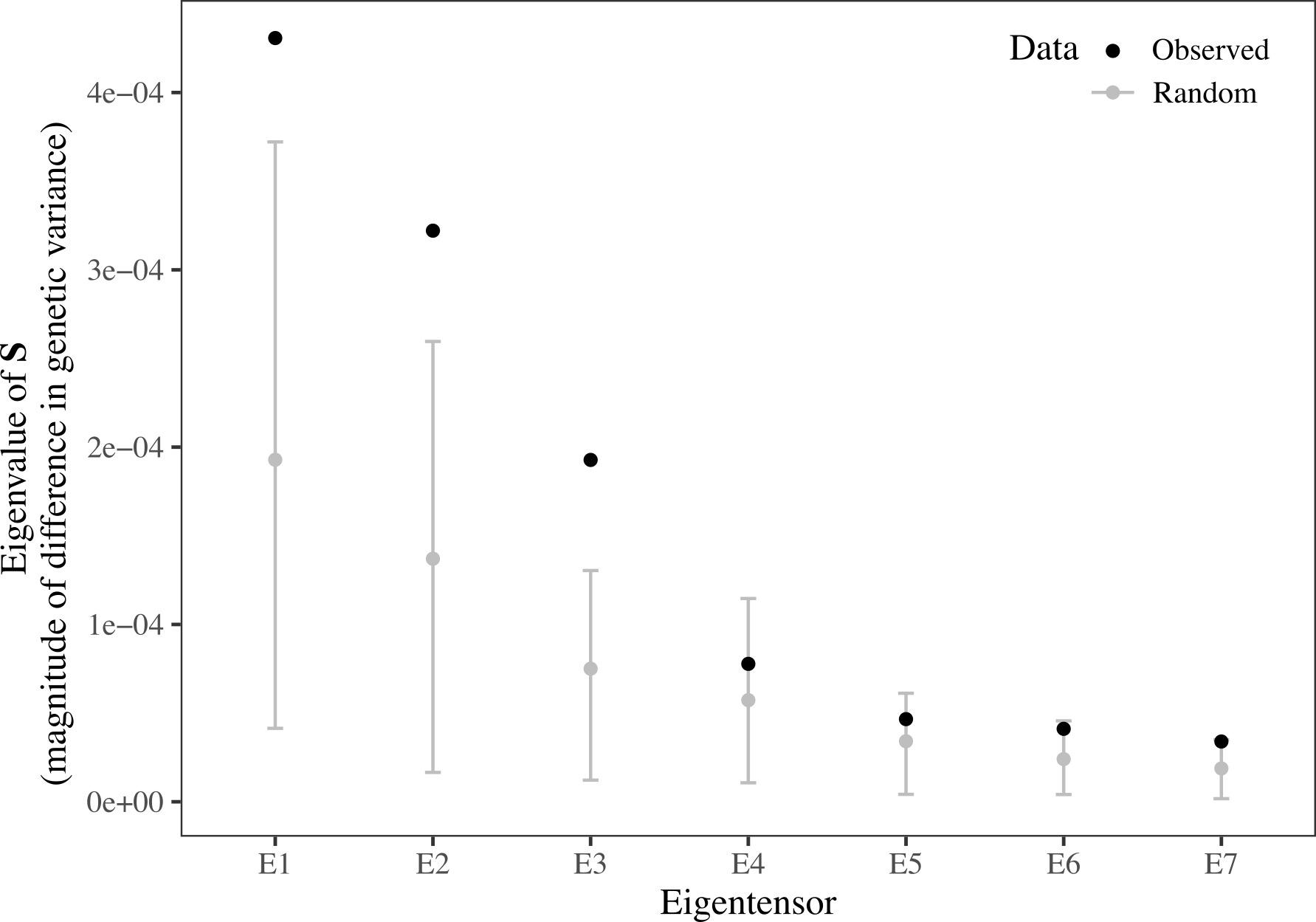

**Fig. S8** Comparing the magnitude of difference in genetic variance captured by the tensor applied to the observed estimates of genetic variance (black circles) versus the null distribution (grey). The grey credible intervals represent the 95% HPD interval for the null distribution, which is calculated by re-applying equation 1 to data containing no differences in genetic variance and taking the mean from each implementation. The first three eigentensors describe greater differences in genetic variance than expected under the null distribution.

**Table S4** Summary of the covariance tensor analysis that captured differences in **G** across elevation for both species. Presented are the 7 non-zero eigentensors. Eigenvalue of **S** represents the amount of difference in genetic variance described by each eigentensor, with ‘Prop. **S**’ representing the proportion of the total difference in genetic variance. λ represents the amount of difference in genetic variance described by each eigenvector of eigentensor, with ‘Prop λ’ representing the proportion that each eigenvector contributes to describing the difference in genetic variance described by the corresponding eigentensor. HPD represents the 90% Highest Posterior Density intervals.

| **Eigen-tensor** | **Eigenvalue of S (HPD)** | **Prop. S** | **Eigenvector of eigentensor** | **Prop.** λ | λ | **P2A** | **IW** | **Nind** | **SLA** | **Flav** |
| --- | --- | --- | --- | --- | --- | --- | --- | --- | --- | --- |
| E1 | 0.000431  (0.000142, 0.000729) | 0.35 | ***e***_1,1_ | 0.66 | -0.95 | 0.40 | 0.47 | -0.46 | 0.05 | -0.63 |
|  |  |  | ***e***_1,2_ | 0.19 | -0.27 | 0.91 | -0.22 | 0.28 | -0.12 | 0.19 |
|  |  |  | ***e***_1,3_ | 0.12 | -0.17 | -0.08 | -0.51 | 0.41 | 0.19 | -0.72 |
|  |  |  | ***e***_1,4_ | 0.02 | -0.03 | 0.10 | 0.09 | 0.03 | 0.97 | 0.19 |
|  |  |  | ***e***_1,5_ | 0.01 | 0.01 | 0.06 | -0.68 | -0.73 | 0.07 | 0.07 |
| E2 | 0.000322  (6.33e-05, 0.000598) | 0.27 | ***e***_2,1_ | 0.61 | 0.89 | 0.96 | -0.15 | 0.00 | -0.03 | -0.23 |
|  |  |  | ***e***_2,2_ | 0.30 | -0.44 | -0.03 | 0.57 | -0.62 | 0.13 | -0.53 |
|  |  |  | ***e***_2,3_ | 0.04 | -0.06 | -0.19 | -0.25 | 0.08 | -0.78 | -0.54 |
|  |  |  | ***e***_2,4_ | 0.04 | -0.05 | 0.15 | 0.12 | -0.54 | -0.55 | 0.61 |
|  |  |  | ***e***_2,5_ | 0.01 | 0.01 | 0.13 | 0.76 | 0.57 | -0.27 | 0.08 |
| E3 | 0.000193  (3.3e-05, 0.000332) | 0.16 | ***e***_3,1_ | 0.45 | -0.71 | -0.10 | -0.04 | 0.06 | -0.24 | 0.96 |
|  |  |  | ***e***_3,2_ | 0.43 | 0.69 | -0.35 | -0.69 | 0.62 | 0.11 | -0.08 |
|  |  |  | ***e***_3,3_ | 0.07 | 0.11 | 0.82 | -0.53 | -0.14 | 0.13 | 0.11 |
|  |  |  | ***e***_3,4_ | 0.04 | -0.06 | -0.26 | -0.05 | -0.33 | 0.88 | 0.21 |
|  |  |  | ***e***_3,5_ | 0.01 | -0.02 | 0.36 | 0.49 | 0.70 | 0.37 | 0.10 |
| E4 | 7.78e-05  (6.86e-06, 0.000149) | 0.06 | ***e***_4,1_ | 0.63 | 0.90 | 0.63 | -0.54 | 0.55 | -0.04 | 0.13 |
|  |  |  | ***e***_4,2_ | 0.30 | -0.43 | 0.77 | 0.40 | -0.46 | -0.08 | -0.15 |
|  |  |  | ***e***_4,3_ | 0.04 | -0.06 | -0.05 | 0.08 | -0.06 | -0.82 | 0.57 |
|  |  |  | ***e***_4,4_ | 0.02 | 0.03 | -0.08 | -0.48 | -0.26 | -0.49 | -0.67 |
|  |  |  | ***e***_4,5_ | 0.01 | 0.01 | 0.01 | -0.56 | -0.65 | 0.29 | 0.43 |
| E5 | 4.66e-05  (4.65e-06, 9.02e-05) | 0.04 | ***e***_5,1_ | 0.59 | -0.92 | 0.17 | 0.41 | -0.40 | -0.66 | 0.46 |
|  |  |  | ***e***_5,2_ | 0.17 | -0.27 | 0.18 | 0.00 | -0.01 | 0.60 | 0.78 |
|  |  |  | ***e***_5,3_ | 0.17 | 0.26 | 0.38 | 0.44 | -0.56 | 0.42 | -0.41 |
|  |  |  | ***e***_5,4_ | 0.05 | -0.07 | 0.77 | 0.14 | 0.60 | -0.13 | -0.07 |
|  |  |  | ***e***_5,5_ | 0.03 | 0.04 | 0.44 | -0.79 | -0.41 | -0.13 | -0.01 |
| E6 | 4.11e-05  (5.61e-06, 7.56e-05) | 0.03 | ***e***_6,1_ | 0.45 | 0.76 | 0.78 | -0.05 | 0.10 | -0.32 | 0.53 |
|  |  |  | ***e***_6,2_ | 0.36 | -0.61 | 0.45 | -0.49 | 0.31 | 0.15 | -0.66 |
|  |  |  | ***e***_6,3_ | 0.12 | 0.21 | -0.29 | -0.18 | 0.19 | -0.90 | -0.18 |
|  |  |  | ***e***_6,4_ | 0.06 | -0.10 | 0.33 | 0.75 | -0.23 | -0.20 | -0.49 |
|  |  |  | ***e***_6,5_ | 0.02 | 0.03 | 0.09 | -0.41 | -0.89 | -0.12 | -0.09 |
| E7 | 3.4e-05  (1.9e-06,  6.6e-05) | 0.03 | ***e***_7,1_ | 0.45 | -0.78 | 0.29 | 0.29 | -0.30 | 0.82 | -0.28 |
|  |  |  | ***e***_7,2_ | 0.33 | 0.56 | 0.15 | 0.32 | -0.34 | -0.53 | -0.69 |
|  |  |  | ***e***_7,3_ | 0.16 | -0.27 | -0.12 | 0.62 | -0.46 | -0.14 | 0.61 |
|  |  |  | ***e***_7,4_ | 0.04 | 0.07 | 0.72 | 0.37 | 0.56 | -0.13 | 0.16 |
|  |  |  | ***e***_7,5_ | 0.03 | -0.04 | 0.60 | -0.53 | -0.52 | -0.13 | 0.24 |

**Table S5** Estimates of selection coefficients for the phenotypic selection gradients (***β***) estimated for *S. aethnensis* at each elevation away from its home site.

| Elevation | Fixed effect | Coefficient | Df | Sum of Squares | Pr(>Chi) | 95% Confidence interval |
| --- | --- | --- | --- | --- | --- | --- |
| **500m** | Intercept | -3.1 |  |  |  | -6.95, 0.75 |
|  | **Complexity** | **2.9** | **1** | **132.463** | **<0.00001** | **1.93, 3.88** |
|  | Indent width | 0.16 | 1 | 0.096 | 0.87420 | -1.81, 2.13 |
|  | Number indents | 0 | 1 | 0 | 0.99904 | -1.73, 1.74 |
|  | SLA | -0.21 | 1 | 2.366 | 0.43139 | -0.72, 0.31 |
|  | **Flavonol** | **1.24** | **1** | **41.496** | **0.00103** | **0.5, 1.98** |
| **1,000m** | Intercept | 0.13 |  |  |  | -0.78, 1.05 |
|  | **Complexity** | **0.33** | **1** | **1.709** | **0.00536** | **0.1, 0.56** |
|  | Indent width | -0.1 | 1 | 0.04 | 0.66773 | -0.54, 0.35 |
|  | Number indents | 0.1 | 1 | 0.051 | 0.62903 | -0.31, 0.51 |
|  | **SLA** | **0.43** | **1** | **2.996** | **0.00023** | **0.2, 0.66** |
|  | Flavonol | 0.1 | 1 | 0.351 | 0.20591 | -0.06, 0.26 |
| **1,500m** | Intercept | 0.34 |  |  |  | -0.92, 1.59 |
|  | Complexity | 0.15 | 1 | 0.191 | 0.28639 | -0.13, 0.43 |
|  | Indent width | -0.05 | 1 | 0.005 | 0.86020 | -0.67, 0.56 |
|  | Number indents | 0.03 | 1 | 0.002 | 0.92000 | -0.57, 0.63 |
|  | **SLA** | **0.17** | **1** | **1.28** | **0.00597** | **0.05, 0.29** |
|  | **Flavonol** | **0.37** | **1** | **1.995** | **0.00062** | **0.16, 0.58** |

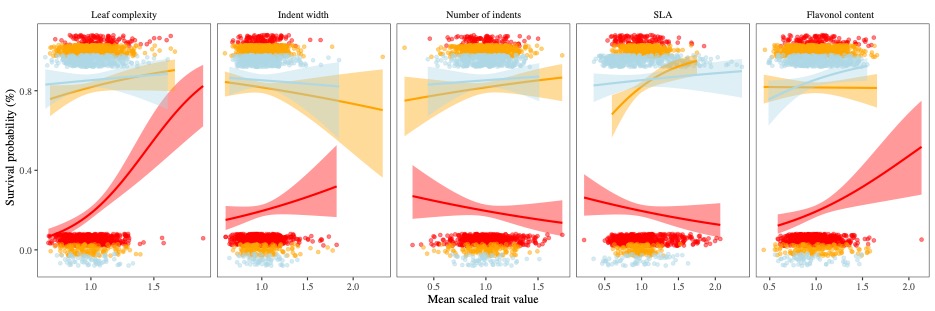

**Fig. S9** Visualising the selection gradients calculated (from **Table S5**) for the five leaf traits (each panel), only using data collected for *S. aethnensis* at 500-1,500m. Regressions are calculated using survival as a binary trait with the shaded ribbons representing one standard error.

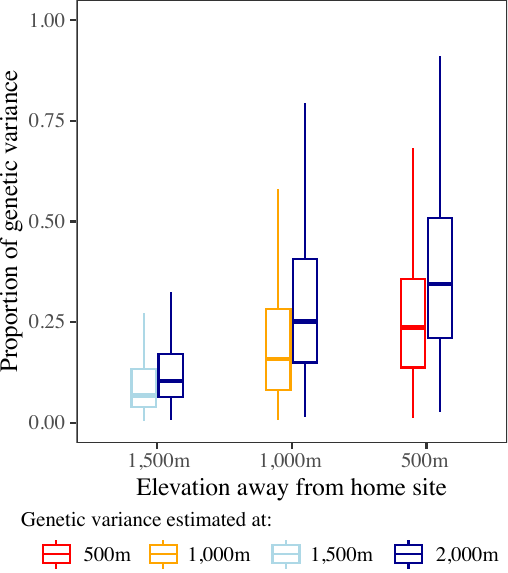

**Fig. S10** Projection of genotypic selection gradients, which were calculated with the same traits and used the mean of each full-sibling family to quantify genotypic selection on each trait. Results are similar to those presented in text for the phenotypic selection gradients.

**Methods S1**
To visualise how the two species differed across elevation we first constructed a D-matrix, the covariance matrix representing differences in mean multivariate phenotype between species and across elevation. To construct **D**, from the MANOVA we extracted the Sums of Squares and Cross-Product (SSCP) matrices for each fixed effect (SSCP_S_ = species; SSCP_E_ = elevation; SSCP_S×E_ = species×elevation) and the error term (SSCP_R_). We then estimated SSCP_H_ ($\mathrm{SSCP}_{H}=\mathrm{SSCP}_{S}+\mathrm{SSCP}_{E}+\mathrm{SSCP}_{S\times E}$), which calculates the difference in mean across all elevations for both species. We calculated Mean Square (MS) matrices by dividing the SSCP matrices by their corresponding degrees of freedom ($\mathrm{MS}_{H}=\frac{\mathrm{SSCP}_{H}}{7};\mathrm{MS}_{R}=\frac{\mathrm{SSCP}_{R}}{6,446}$). We then estimated **D** using

$\mathbf{D}=\frac{\mathrm{MS}_{H}-\mathrm{MS}_{R}}{nf} ,$	*(1)*

where *nf* represents the average number of individuals measured for each species at each elevation, calculated from equation 9 in Martin et al. (2008). We used the eigenvectors of **D** to visualise differences in multivariate phenotype across elevation for both species.
